# Gene Correction for SCID-X1 in Long-Term Hematopoietic Stem Cells

**DOI:** 10.1101/397463

**Authors:** Mara Pavel-Dinu, Volker Wiebking, Beruh T. Dejene, Waracharee Srifa, Sruthi Mantri, Carmencita E. Nicolas, Ciaran Lee, Gang Bao, Eric J. Kildebeck, Niraj Punjya, Camille Sindhu, Matthew A. Inlay, Nivi Saxena, Suk See DeRavin, Harry Malech, Maria Grazia Roncarolo, Kenneth I. Weinberg, Matthew H. Porteus

## Abstract

Gene correction in human long-term hematopoietic stem cells (LT-HSCs) could be an effective therapy for monogenic diseases of the blood and immune system. High frequencies of reproducible targeted integration of a wild-type cDNA into the endogenous start codon of a gene in LT-HSCs could provide a robust genome editing approach to cure genetic diseases in which patients have different mutations throughout the gene. We describe a clinically relevant method for correcting X-linked severe combined immunodeficiency (SCID-X1). By using a highly specific and active CRISPR/Cas9-AAV6 based strategy and selection-free approach, we achieve up to 20% genome integration frequencies in LT-HSCs of a full-length *IL2RG* cDNA at the endogenous start site as demonstrated by serial transplantation and analysis of genome edited human cells eight months following initial transplantation. In addition to high frequencies of functional gene correction in LT-HSCs we observed no evidence of abnormal hematopoiesis following transplantation, a functional measure of the lack of genotoxicity. Deep analysis of potential off-target activity detected two sites with low frequency (<0.3%) of off-target mutations. The level of off-target mutations was reduced to below the limit of detection using a high fidelity Cas9. Moreover, karyotype evaluation identified no genomic instability events. We achieved high levels of genome targeting frequencies (median 45%) in CD34^+^ HSPCs from six SCID-X1 patients and demonstrate rescue of lymphopoietic defect of patient derived cells both *in vitro* and *in vivo.* In sum, our study provides specificity, toxicity and efficacy data supportive of clinical development of genome editing to treat SCID-Xl.

---

X-linked severe combined immunodeficiency (SCID-X1) is a rare, primary immune deficiency (PID) caused by mutations in the *IL2RG* gene on the X-chromosome. The gene encodes a shared subunit of the receptors for IL-2, IL-4, IL-7, IL-9, IL-15, and IL-21. Without early treatment, affected male infants die in the first year of life from infections. Although allogeneic hematopoietic cell transplant (allo-HCT) is considered the standard of care for SCID-X1, it holds significant risks due to potential incomplete immune reconstitution, graft versus host disease (GvHD) and a decreased survival rate in the absence of an HLA-matched sibling donor^1^. Importantly, because of the selective advantage of lymphoid progenitors expressing normal *IL2RG*, only a small number of genetically corrected hematopoietic stem and progenitor cells (HSPCs) are needed to reconstitute T-cell immunity. The selective advantage has been demonstrated both by allo-HCT and by rare experiments of nature, in which a spontaneous reversion in a hematopoietic progenitor gives rise to a functional T-cell repertoire in a SCID-X1 patient^2,3^. The rare patients with reversion mutations and patients who received allo-HCT without giving conditioning chemotherapy also highlights the importance of achieving gene correction in long-term hematopoietic stem cells (LT-HSCs) to achieve sustained clinical benefit as patients who do not have corrected LT-HSC engraftment often end up losing a robust and functional T-cell immune system over time.

Gene therapy is an alternative therapy to allo-HSCT. Using integrating viral vectors, such as gamma-retroviral and lentiviral vectors, extra copies of a functional *IL2RG* gene are semi-randomly integrated into the genome of SCID-X1 patient derived CD34^+^ hematopoietic stem and progenitor cells (HSPCs). This strategy has resulted in both successes and setbacks. While most patients treated with 1^st^ generation of gene therapy survived and benefited from the therapy, a substantial fraction (>25%) of patients developed leukemia from insertional oncogenesis^*4, 5, 6*^. It is concerning that patients developed leukemia from insertional oncogenesis both early and late, 15 years after transplantation of retroviral-based engineered cells (Six, E., et al., Abstract #753, ASGCT Meeting, 2017). Constitutive activation of the transgene^*7*^, the choice of vectors^*8*^ and specific details of the gene therapy procedure have all been proposed as factors contributing to the risk of leukemia and myelodysplastic syndrome that occurred in several trials for primary immunodeficiency disorders (PIDs) including SCID-X1^*9, 10*^, chronic granulomatous disease (CGD)^*11, 12*^ and Wiskott-Aldrich Syndrome (WAS)^*13*^. With 2^nd^ generation self-inactivating (SIN) vectors, multiple SCID-X1 patients have successfully reconstituted T-cell immunity in the absence of early leukemic events^*14, 15, 16*^ with a follow up of up to 7 years. However, the follow-up of these therapies remains too short to assess the long-term genotoxicity risk of the newer generation vectors, as transformation of T-cells growth can take more than 10 years to manifest (Six, E., et al., Abstract #753, ASGCT Meeting, 2017).

One strategy to convert the semi-random delivery and integration of a corrective *IL2RG* cDNA into the genome is through a genome editing (GE) approach. GE is a means to alter the DNA sequence of a cell, including somatic stem cells, with nucleotide precision. Using homologous recombination mediated genome editing (HR-GE), the approach can target a cDNA transgene into its endogenous locus, thereby preserving normal copy number and upstream and downstream non-coding elements that regulate expression^*17, 18, 19*^. Currently the highest frequencies of GE can be achieved by using an engineered nuclease to create a site-specific double-strand breaks (DSBs) in the cell’s genomic DNA^*20,21*^. When the DSB is repaired by non-homologous end joining (NHEJ), small insertions and deletions (INDELs) can be created at a specific genomic target site - a powerful approach to inactivating genetic elements but not generally useful for correcting mutant genes^*22,23*^. In contrast, when the DSB is repaired by either homologous recombination (using a classic gene targeting donor vector) or by single-stranded template repair (using a single-stranded oligonucleotide (ssODN)), precise sequence changes can be introduced. This provides a method to precisely change DNA variants, including converting disease-causing mutations into those that do not^*24*^.

Among the multiple GE platforms that use artificial nucleases to generate DSBs ^*25, 26, 27, 17, 28*^ CRISPR/Cas9 system has accelerated the field of GE because of its ease of use and high activity in a wide variety of cells, including somatic stem cells. When CRISPR/Cas9 is delivered into primary human cells, including human CD34^+^ HSPCs as a ribonucleoprotein (RNP) complex using fully synthesized single-guide RNA molecules (sgRNAs) with end modifications to protect the guide from exonuclease degradation, high frequencies of INDELs are achieved^*29*^. Moreover, when the delivery of an RNP complex is combined with delivery of the gene targeting donor molecule in a recombinant AAV6 (rAAV6) viral vector, high frequencies of homologous mediated editing in human HSPCs are obtained^*24*^. The use of rAAV6 donor vectors have been successfully used with other nuclease systems as well, including zinc finger nucleases (ZFNs) and in other cell types, such as primary human T-cells^*18, 30, 31*^. Therefore, this HR-GE approach could transform the random nature of current gene therapy design to a controlled, precise and safer treatment option. By using AAV6 as a classic gene targeting donor a full cDNA can be introduced at the endogenous target, an event that cannot be achieved using a ssODN because ssODN mediated HDR cannot integrate full genes.

The central questions that are currently being investigated in the field of genome editing are focused on the ability to efficiently deliver the genome editing reagents to clinically relevant cell types, attaining functional level of expression from the codon optimized cDNA, preserving stem cell potential of edited cells, and establishing lack of genotoxicity derived from genome editing methodology.

Here, we describe a clinically relevant, selection-free “universal” CRISPR/Cas9-rAAV6 GE methodology that could potentially correct >97% of known *IL2RG* pathogenic mutations. We call this approach “functional gene correction” because it is not directly correcting a mutation but instead is doing so by using the targeted integration of cDNA to functionally correct downstream mutations. The ∼2-3% of patients with deletions of the gene that could not be functionally corrected using this strategy. We demonstrate that a functional, codon optimized *IL2RG* cDNA can be precisely and efficiently integrated at the endogenous translational start site in CD34^+^ HSPCs of healthy male donors (HD, n=13) or SCID-X1 patients (n=6) at comparable frequencies (median HR = 45%). We show the process is highly reproducible in both peripheral blood derived and umbilical cord blood derived CD34^+^ HSPCs. We further demonstrate the functionality of the full-length codon optimized *IL2RG* cDNA by showing that T-cells with the cDNA knock-in retain normal proliferation and signaling response to cytokines. Using transplantation into immunodeficient (NSG) mice we show that process is both effective (with functional correction of 10-20% of LT-HSCs) and safe (no evidence of abnormal hematopoiesis). These *in vivo* functional results are based on transplantation of ∼21 million *IL2RG* targeted healthy donor CD34^+^ HSPCs (combined primary and secondary engraftment studies) and ∼7 million *IL2RG* targeted SCID-Xl-HSPCs. By achieving a median of 45% human engraftment from functionally corrected cells, a multi-lineage differentiation *in vivo* from primary human engraftment and preservation of up to 20% human engraftment of *IL2RG* cDNA genome targeted HSPCs at 8 months following secondary human engraftment, we proved that clinically relevant correction levels were achieved in CD34^+^ LT-HSCs. These results match and exceed the predicted therapeutic threshold determined through a mouse model^18^. Lastly, we show no evidence of significant genotoxicity as demonstrated by Next Generation Sequencing (NGS) and karyotype analysis. Together, this study establishes a pre-clinical proof-of-concept for a safe, precise and highly efficient genome editing strategy that holds the potential of a curative therapy for SCID-X1.

## Results

### Establishing a universal gene correction strategy for *IL2RG* locus in CD34^+^ HSPCs

SCID-X1 is caused by pathogenic mutations spanning the entire *IL2RG* open reading frame. Therefore, we developed a gene targeting strategy by integrating a complete cDNA at the endogenous *IL2RG* translational start site **(Fig 1a, central panel)** that would correct the vast majority (∼97%) of known SCID-X1 pathogenic mutations and ensure regulated endogenous expression in CD34^+^ HSPCs derived progeny. By achieving efficient integration frequencies in the genome of CD34^+^ LT-HSCs, our approach could ensure life-long therapeutic benefits for the patient **(Fig 1a, right schematic)**.

**Figure 1.**
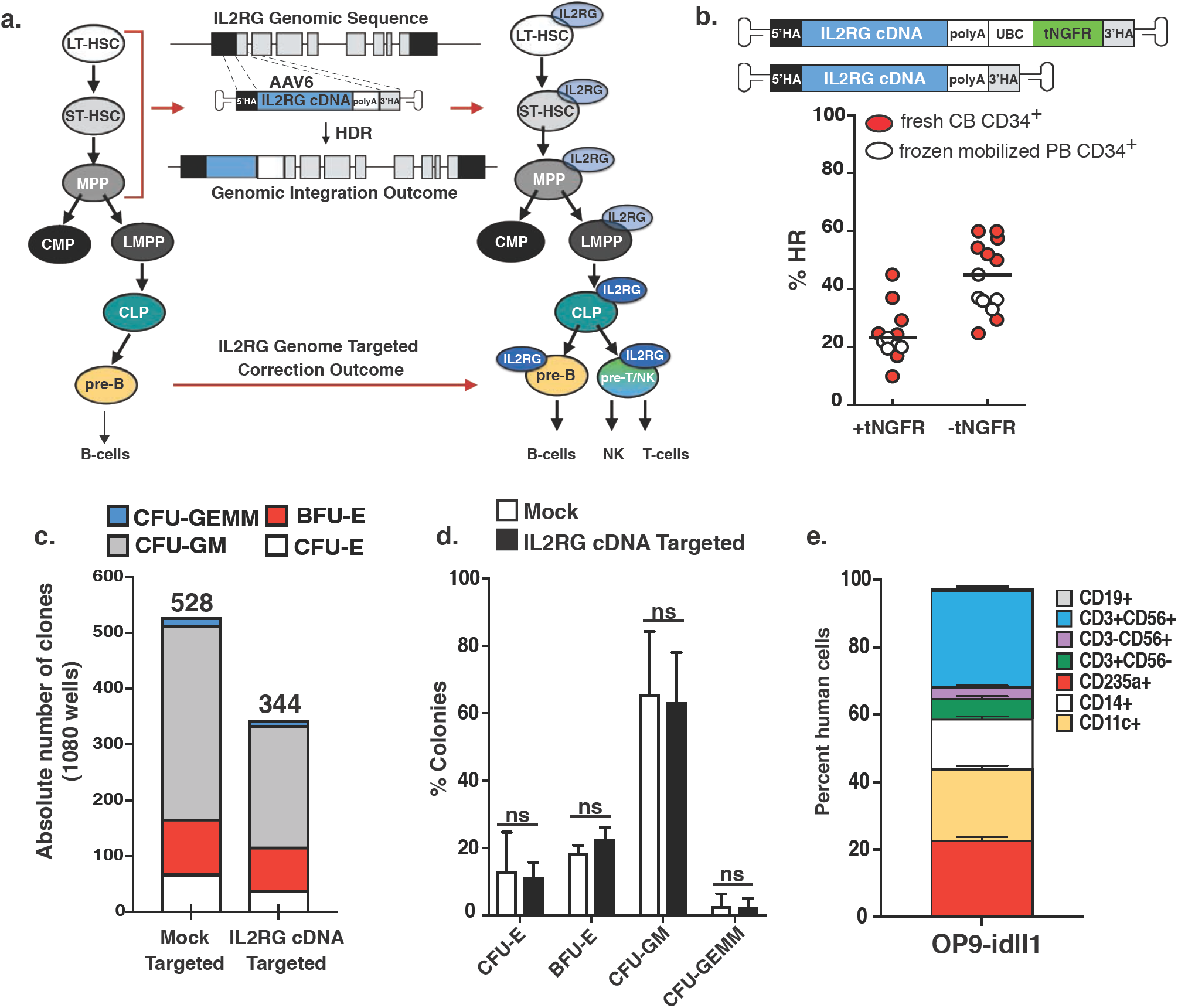
In vitro validation of medium-scale CRISPR/Cas9-AAV6 based IL2RG cDNA targeting in CD34^+^ HSPCs. **a.** Overview of genomic targeting at IL2RG locus using a full length, codon optimized and promoter-less IL2RG cDNA. Schematic of genomic integration and genomic correction outcomes are shown. Top: Schematic of IL2RG corrective donors containg selectable marker (+tNGFR) or not (-tNGFR). Bottom: Day 4, medium-scale (1.0 × 10^6^) ex-vivo IL2RG cDNA targeting frequencies of frozen mobilized peripheral blood CD34^+^ HSPCs (white circles) or freshly purified cord blood male-derived CD34^+^ HSCPs (red circles). absolute targeting frequencies measured by ddPCR. Bar is median: 23.2% (+tNGFR, n=11), 45% (-tNGFR, n=13) Single cell based methylcellulose assay using mock targeted (nucleofected only) or IL2RG cDNA targeted (-tNGFR donor) CD34^+^ HSPCs. Absolute number of clones derived from each targeting condition are shown. Fraction of the total for each type of colony scored. Bars (mean ± s.e.m.), ns = not specific (Welch’s t-test) Gene correction of SCID-X1 derived CD34^+^ HSPCs and multilineage differentiation using OP9-idll1 in vitro system. Shown are lineages derived from IL2RG cDNA targeted HSPCs. SCID-X1 patient 2 is shown with 45% gene targeted frequency at day 2 pre-sorting onto OP9-idll1 feeders. No growth was derived from uncorrected CD34^+^ cells. LT-HSPCs, long-term hematopoietic stem cells; ST-HSC, short term hematopoietic stem cells; MPP, multi-potent progenitor; CMP, common myeloid progenitor; LMPP, lymphoid multi-potent progenitor; CLP, common lymphoid progenitor; HSPCs, hematopoietic stem and progenitor cells. Bars (mean ± s.e.m.)

Towards this goal, we screened seven different sgRNA chemically modified guides^29^ for activity in exon 1 of the *IL2RG* gene **(Supplemental Fig 1a)** and selected sg-1, previously described^29^, as the best candidate because of the location of the double-strand break (DSB) it creates (one nucleotide downstream from the translational start site), on-target INDEL frequencies (92.9% ± 0.6, mean ± *s.e.m*) **(Supplementary Fig 1b)** and for high cellular viability > 80% **(Supplementary Fig 1c).** We optimized the process using this gRNA and found that a truncated guide^32^ of 19 nucleotides (19nt) gave >90% INDEL frequencies (equivalent to the full 20nt gRNA) while maintaining high cellular viability (**Supplemental Figures 1, 2, and 3).** We found a significant decrease in on-target editing using truncated 18nt and 17nt gRNAs **(Supplementary Fig 1d)**. Next Generation Sequencing (NGS) **(Supplementary Fig. 1e)** further corroborated the INDELs obtained by TIDE analysis, results which are in line with a recent study that demonstrated that TIDE analysis is as good as NGS in terms of measuring the percentage of INDELS and the distribution in a large population of cells^33^. Throughout the rest of the studies we used the 19nt gRNA at a medium scale process (1 million cells per electroporation).

**Figure 2.**
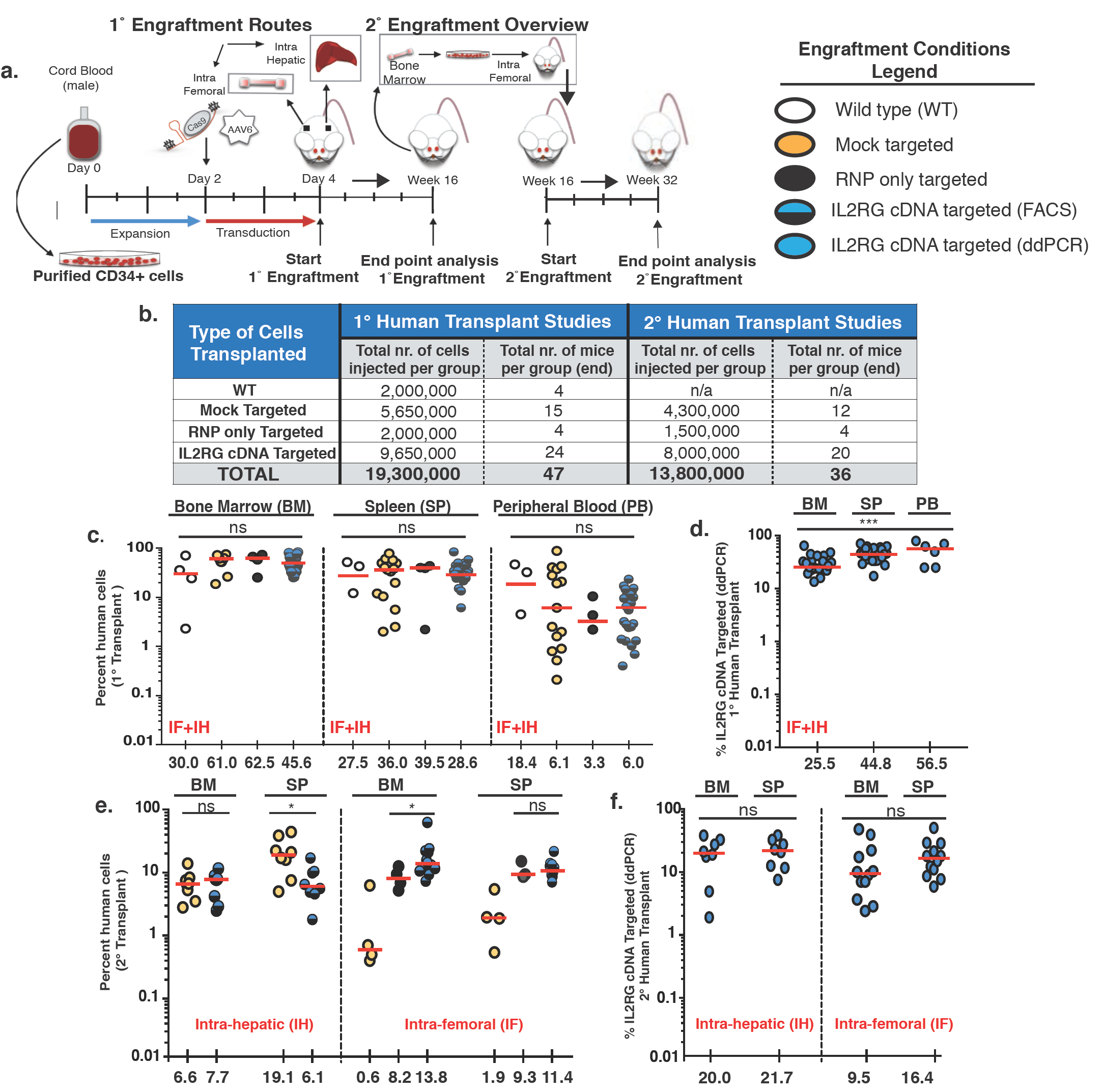
In vivo normal hematopoietic reconstitution from medium-scale IL2RG cDNA targeted CD34^+^ HSPCs from healthy donors. **a.** Schematic representation of primary and secondary transplants into sub-lethaly irradiated NSG mice. Adult mice transplanted intra-femoral with either WT CD34^+^ HSPCs (white circles) or mock targeted (yellow circles) or RNP only (black circles) or un-selected IL2RG cDNA targeted (blue-black circles) HSPCs. 3-4 days old NSG pups transplanted intra-hepatic with either mock or IL2RG targeted HSPCs. Summary of total number of cells and mice injected per condition for primary (1°) and secondary (2°) transplants. **c.** Combined intra-femoral (IF) and intra-hepatic (IH) human cells engraftment (hCD45^+^ HLA A-B-C^+^) 16 weeks after transplant into indicated organs. BM, bone marrow, SP, spleen, PB, peripheral blood. **d.** %IL2RG cDNA targeted HSPCs within human graft in indicated organs, as assesed by ddPCR. BM (n=24), SP (n=24), PB (n=6) (\*\*\**P*-value =0.0008, one-way ANOVA) **e.** Percent human engraftment in indicated organs as in (**c**) 16 weeks post 2 transplants of human CD34^+^ from mice. \**P*-value SP-IH = 0.025, \**P*-value BM-IF = 0.043 (Welch’s t-test). **f.** %IL2RG targeted HSPCs quantified by ddPCR 32 weeks after engraftment. Median shown. ns, not significant. ddPCR, digital drop PCR.

**Figure 3.**
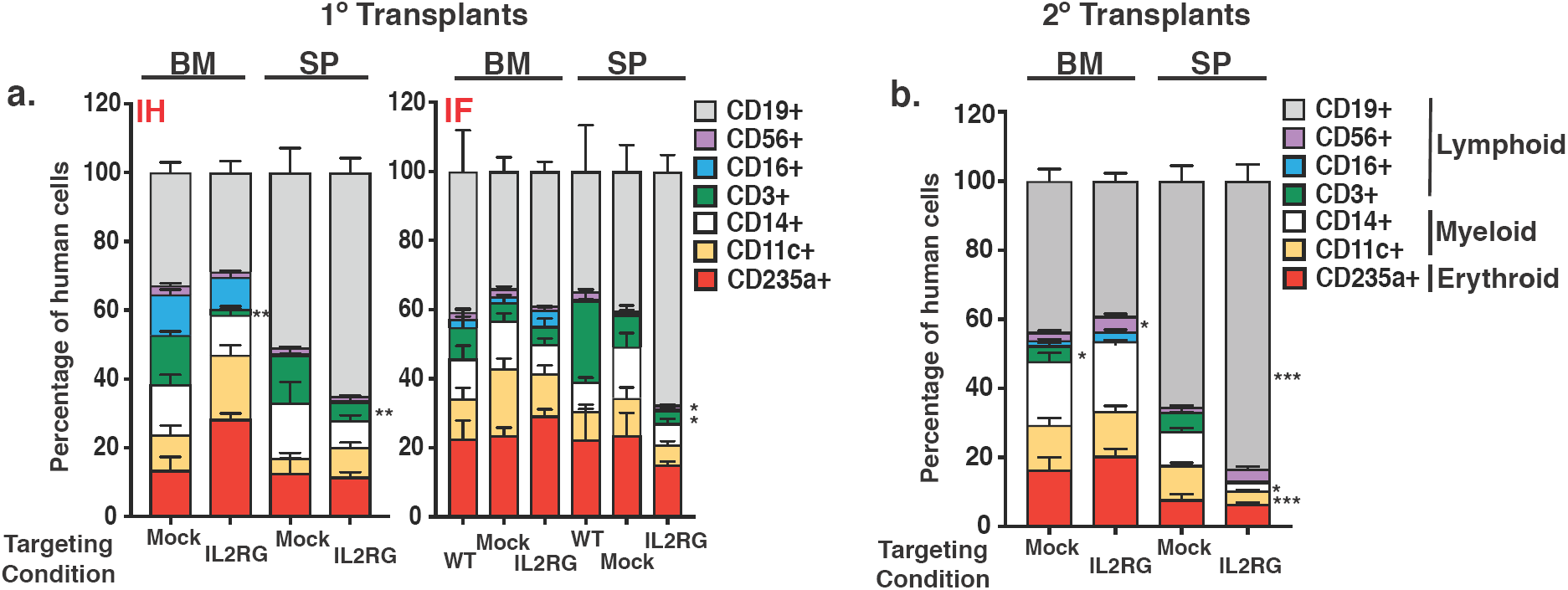
Normal multilineage development and validation of IL2RG cDNA targeting in the long-term (LT) HSC population. **a.** Percent cellular composition of the lymphoid, myeloid and erythroid lineage derived from 1° human engraftment, shown in indicated organs and targeting conditions. Left panel: intra-hepatic (IH) transplant analysis.CD3^+^ BM: \*\**P*-value = 0.0017, CD3^+^ SP: \*\**P*-value = 0.007 (Welch’s t-test) Right panel: intra-femoral (IF) transplant analysis. CD3^+^ SP: \**P*-value = 0.023, CD56^+^ BM: \**P*-value = 0.015 (Kruskal-Wallis test). **b.** Same as (**a**) derived from secondary transplants. Data shown is combined intra-hepatic and intra-femoral primary transplants. CD3^+^ BM: \**P*-value = 0.015, CD56^+^ \**P*-value = 0.025, CD19^+^ SP: \*\*\**P*-value = 0.0002, CD14^+^ SP: \**P*-value = 0.0112, CD11c^+^ SP: \*\*\**P*-value = 0.0004, Bars: mean ± *s.e.m*

We designed a codon optimized *IL2RG* cDNA functional correction donor with homology arms centered on the sg-1 guide sequence **(Supplemental Fig 4a-b).** A functional correction cDNA donor was cloned into an AAV6 vector, with or without the presence of a selectable marker cassette (the truncated nerve growth factor receptor (tNGFR) under the control of the human Ubiquitin C (UBC) promoter) **(Fig 1b, top panel).** The efficiency of genome-targeting integration was determined in both frozen mobilized-peripheral blood (mPB) and in freshly isolated cord blood (CB) derived-CD34^+^ HSPCs from healthy male donors **(Fig 1b, bottom panel)**. We observed a median gene-targeting frequency of 23.2% (range 9.9% – 45.0%) for the +tNGFR donor and 45.0% (range 24.7% - 60.0%) for the -tNGFR *IL2RG* donor **(Fig. 1b, bottom panel)**, as measured by a Digital Droplet PCR (ddPCR) *IL2RG* specific assay **(Supplemental Fig. 5a-f)**. Since the selection free cassette gave such high frequencies of targeted integration we reasoned that the need for a selection marker was not needed because of the selective advantage of functionally corrected lymphoid cells and their progenitors have and would entail a simpler clinical cell manufacturing process.

To determine the myeloerythroid differentiation potential of *IL2RG* cDNA genome targeted CD34^+^ HSPCs we performed methylcellulose clonal assays. After Cas9/gRNA-rAAV6 based *IL2RG* cDNA targeting, HSPCs were single cell plated in a 96-well methylcellulose plates with growth factors and scored for colony formation at day 14. Although the number of colonies was reduced by ∼35% in *IL2RG* cDNA targeted samples compared to mock-targeted HSPCs (where neither the sgRNA nor the donor were introduced) **(Fig. 1c)**, the distribution of types of colony-forming units (CFU) was the same from *IL2RG* cDNA targeted HSPCs and mock-targeted HSPCs (where neither the sgRNA nor the donor were introduced) **(Fig. 1d)**. Furthermore, both targeting conditions gave rise to erythroid (CFU-E, BFU-E), granulocytes/macrophages (CFU-GM) and multi-lineage (CFU-GEMM) colonies, without lineage skewing. Genotyping of colonies confirmed that *IL2RG* cDNA targeted derived-colonies (n = 344) showed an overall targeting frequency of 45.7% ± 2.44 (mean ± *s.e.m.*) **(Supplemental Fig 6)** (a frequency equivalent to that determined by ddPCR from the bulk population of targeted HSPCs) **(Fig. 1b, bottom panel –tNGFR).** Of note, bi-allelic modification is not relevant as the cells were derived from male donors and have a single X-chromosome. In sum, the *in vitro* differentiation assay of targeted *IL2RG* cDNA CD34^+^ HSPCs demonstrated no perturbation of the myeloerythroid differentiation potential and provided evidence against significant toxicity derived from genome edited cells.

To assess the hematopoietic differentiation potential of the codon optimized *IL2RG* cDNA donor, we used the OP9-idll1 stromal cell line *in vitro* system. In this system a lentiviral vector confers the doxycycline (DOX) inducible expression of the Notch ligand Dll1, as previously described^34^. In the presence of a cocktail of cytokines permissive for myeloerythroid and lymphoid differentiation, multipotent human CD34^+^ HSPCs will generate only myeloerythroid and B cell lineage before induction of dll1 expression, but becomes permissive for T and NK cell generation in the same well after addition of DOX to induce dll1 expression. CD34^+^ HSPCs derived from frozen mPB of SCID-X1 patient (delA;M145fs – patient 2) were gene targeted (functionally corrected) using the CRISPR/Cas9-AAV6 platform. 48 hours following AAV6 transduction, *IL2RG* cDNA targeted or mutant HSPCs were plated (300 cells per well) in a 96-well plate containing OP9-idll1 stromal cells. The total number of cells per well derived from the *IL2RG* cDNA targeted cells was markedly increased, compared to that of mutant cells, indicating a growth dependence on functional IL-7 and IL-15 receptors, for which *IL2RG* is an essential subunit ^*35*^. Following DOX mediated dll1 expression no further growth of mutant CD34^+^ HSPCs was detected on OP9-idll1 stromal cells. In contrast, the *IL2RG* cDNA targeted cells continued to expand in myeloerythroid compartment in addition to the development of B (CD19^+^), T (CD3^+^CD56^−^), NK (CD3^−^CD56^+^) and TNK (CD3^+^CD56^+^) progeny **(Fig. 1e, Supplemental Fig. 7S)**. The OP9-idll1 *in vitro* system is known to generate more CD3^+^CD4^+^ over CD3^+^CD8^+^ cells expressing cell surface markers^36^. We do note that a CD3^+^CD56^+^ (TNK cell) population was generated from our genome corrected SCID-X1 patient derived CD34^+^ HSPCs. This population was reported^37^ to be a lymphoid population that has a non-antibody-dependent cell cytotoxicity-mediated anti-viral activity. CD3^+^CD56^+^ (TNK) cells were derived from CD3^+^CD56^−^ precursors and not from CD3^−^ CD56^+^ or expended from a rare population of CD3^+^CD56^+^. Therefore, the presence of TNK population in our genome corrected SCID-X1 derived CD34^+^HSPCs further demonstrates the range for lymphoid lineage that can arise following *ex-vivo* gene editing correction for IL2RG locus. This assay, therefore, demonstrates the functional correction of *IL2RG* gene from patient-derived CD34+ HSPCs that is necessary for lymphoid development.

In sum, *in vitro*, we observed high targeting frequencies with a full-length codon optimized *IL2RG* cDNA at the translational start site of the endogenous locus. Functionally corrected healthy donor CD34^+^ HSPCs demonstrated preserved multipotency using methylcellulose colony forming unit assay. Functional gene correction of SCID-X1 patient derived peripheral blood mobilized CD34^+^ HSPCs rescued all lymphoid lineages (T, B, and NK) using an *in vitro* assay for lymphoid development.

### Robust *in vivo* hematopoiesis reconstitution from *IL2RG* cDNA targeted HSPCs

To further assess the toxicity and efficacy of our HR-GE methodology, we compared the *in vivo* engraftment potential and multi-lineage (myeloid, erythroid and lymphoid) hematopoietic reconstitution of *IL2RG* cDNA targeted HSPCs with that of normal HSPCs into NOD-scid-gamma (NSG) mice using the selection-free functional correction donor. Following a total of 3.5 to 4 days of *ex viv*o culturing and genomic manipulation, *IL2RG* cDNA targeted, RNP only (cells nucleofected with ribonucleoprotein complex only), mock targeted (nucleofected in the absence of ribonucleoprotein complex) or wild type (WT) CD34^+^ HSPCs were either transplanted by intra-hepatic (IH) injection into sub-lethally irradiated 3-4 day old NSG pups, or by intra-femoral (IF) injection into 6-8 week old NSG mice. The IH system has previously been shown to be superior for assessment of human lymphopoiesis^38^. An experimental schema is shown **(Fig. 2a, primary engraftment panel)**. For primary engraftment studies, a total of 19.3 million cells, derived from 3 different healthy male cord blood CD34^+^ HSPCs were transplanted into a total of 47 mice **(Fig. 2b)**. The kinetics of primary human engraftment was monitored at weeks 8 and 12 in bone marrow aspirates and peripheral blood samples from mice injected either IH or IF. At week 16, an end point analysis was carried out on total bone marrow (BM), spleen (SP) and peripheral blood (PB) samples. Robust human engraftment levels - as shown by hCD45^+^ HLA-ABC^+^ double positive staining, blue/black circles – were obtained with no statistical difference between the *IL2RG* cDNA targeted and control cells - WT, mock, or RNP **(Fig 2c, Supplemental Fig 8).** Transplanted *IL2RG* targeted HSPCs showed a median human engraftment level of 45% in BM (n=24), 28% in SP (n=24) and 6% in PB (n=24) (**Fig 2c).** Importantly, the targeted integration frequency of the codon optimized *IL2RG* cDNA was 25.5% in BM (n=24), 44.8% in SP (n=24) and 56% in PB (n=6) at week 16 post-engraftment, as quantified by ddPCR, blue circles **(Fig. 2d).** Multi-lineage reconstitution was achieved from both mock and *IL2RG* cDNA targeted cells in both the BM and SP samples of transplanted mice **(Fig. 3a).**

In human cells not targeted with the cDNA correction cassette the frequency of INDELs was >90% as determined both by genotyping and TIDE analysis carried out on the IH engrafted *IL2RG* targeted cells at weeks 8, 12 and 16 **(Supplemental Fig. S9a)**. A detailed clonal analysis of INDELs spectrum demonstrates that the three most common types of INDELs generated by the sg-1 IL2RG guide +1, −11, and −13 generate frameshift events that renders *IL2RG* gene inactive **(Supplemental Fig. S9b)**. In sum, our selection free engraftment strategy of *IL2RG* cDNA targeted CD34^+^ HSPCs derived from healthy male donors demonstrate its ability to give rise to normal hematopoiesis. Given that >90% of the non-gene targeted human cells have inactivating INDELs in the *Il2RG* gene, it is likely that the T and NK cells seen in the mice are derived from gene targeted CD34+ HSPCs. But given the paucity of these cells in the mice, we could not definitively molecularly analyze the population for the genotype of the T and NK cells.

### *IL2RG* cDNA genome targeting of LT-HSCs retains normal hematopoiesis

Long-term maintenance of corrected T-cells will require editing of LT-HSCs population, which can support continuous production of blood cells for the lifetime of the patient. We performed secondary transplantation studies to assess the robustness of our CRISPR/Cas9-AAV6 genome targeting platform in editing the LT-HSCs population. CD34^+^ HSPCs were isolated from total bone marrow of mock or *IL2RG* cDNA targeted HSPCs, derived from either primary IH or IF engraftments at week 16 or later. Following overnight culturing, secondary transplants were carried out in sub-lethally irradiated 6-8 week old NSG mice **(Fig 2a, top right secondary engraftment overview)**. At 16 weeks following the secondary transplant, end point analysis – totaling 32 weeks of engraftment into immunodeficient mice – a median human chimerism level (hCD45/HLA-ABC double positive cells) of *IL2RG* cDNA targeted cells ranged from 7.7% to 13.8% (BM) and 6.1% to 11.4% (SP) **(Fig. 2e)**. The median targeted integration frequencies of the *IL2RG* cDNA donor, quantified by ddPCR was 9.5% or 20% (BM) and 16.4% or 21.7% (SP), depending on the mode of primary human engraftment (IH vs IF) **(Fig. 2f).** FACS plots showing BM human engraftment levels from mice injected with cells derived from both conditions are shown **(Supplemental Fig.10, Supplemental Fig. 11).** Multi-lineage analysis in secondary transplants showed no evidence of abnormal hematopoiesis thus providing functional evidence of the safety of the HR-GE process **(Fig. 3b)**.

A summary of the *IL2RG* cDNA targeted engrafted cells is shown in **Table 1**. We report that 20% and 9.5% of human cells in the bone marrow derived from IH-IF and IF-IF secondary xenotransplantation experiments, respectively, retain the codon optimized *IL2RG* cDNA donor integration, demonstrating a clinically significant level of correction of CD34^+^ LT-HSCs. Moreover, our median frequencies of *IL2RG* cDNA targeted in LT-HSCs significantly exceeds those reported by other groups, notably Genovese *et al*^*19*^, Schiroli, J. *et al*^*18*^ and Dever, D.P., *et al*^*24*^. where the percent of HR-GE cells was less than 5% of the human cells engrafted. While Dever *et al.* used Cas9 RNP in their engraftment studies, the Genovese *et al*^*19*^ and Schiroli *et al*^*18*^. engraftment studies were done with zinc finger nucleases (ZFNs). Our results, therefore, represent the first evidence of high frequencies of HR-GE in LT-HSCs using the CRISPR/Cas9 system. No tumors or abnormal hematopoiesis were observed in any mice that were transplanted with genome-modified cells (RNP or *IL2RG* cDNA targeted). Collectively, our primary and secondary xenotransplantation results validate the robustness, effectiveness and lack of genotoxicity of our *IL2RG* cDNA genome targeting approach and strongly supports its advancement towards clinical translation.

**Table 1.**
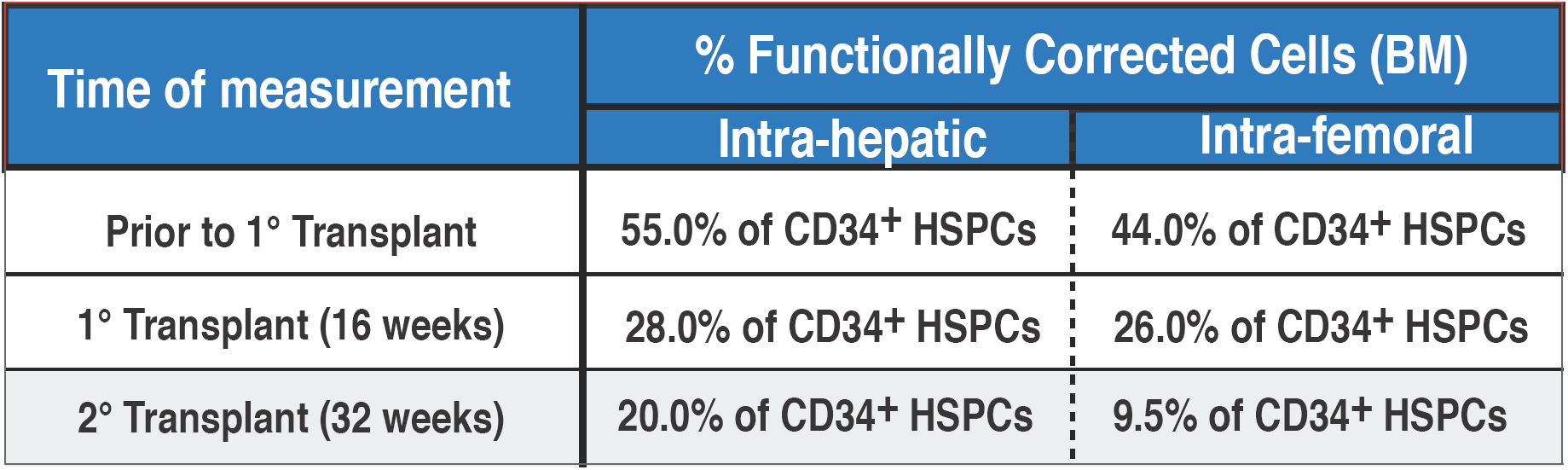
IL2RG cDNA genome targeted frequencies pre- and post-transplant. Percent HDR measured by ddPCR, at the indicated time points, in male derived cord blood CD34^+^ HSPCs targeted with exon 1 IL2RG cDNA. Shown is median. 1° transplant: IH (n = 10), IF (n = 14); 2° transplant: IH (n = 8), IF (n = 12). ddPCR, digital dropplet PCR, IH, intra-hepatic, IF, intra-femoral.

### *In vivo* rescue of lymphopoiesis in *IL2RG* cDNA targeted SCID-X1 HSPCs

We next investigated whether our genome targeting approach was reproducible and efficient in SCID-X1 patient-derived CD34^+^ HSPCs. We edited CD34^+^ HSPCs from six different SCID-X1 patients with a variety of different pathologic mutations (a description of each patients’ pathogenic mutations, gene mapping as well and the source of CD34^+^ derivation are shown) **(Fig. 4a).** Notably five of the six patient’s cells were peripheral blood derived CD34^+^ HSPCs. The *ex-vivo*, CRISPR/Cas9-AAV6 based *IL2RG* cDNA genome targeting platform resulted in greater than 80% viability post-targeting patient-derived cells (n=5) **(Fig. 4b)**. A total of 7.3 million edited CD34+ HSPCs derived from patients 1, 2 and 3 were engrafted into 29 NSG pups **(Fig. 4c)**. Quantification of *IL2RG* cDNA targeting from SCID-X1 patient-derived CD34^+^ HSPCs showed a median of 44.5% (range 30.2% - 47.0%, n=6) **(Fig. 4d)**, a frequency comparable to that obtained from healthy male donors (45%, n=13). Human chimerism was measured at week 16 following IH engraftment, with no statistically significant differences between unmodified and *IL2RG* cDNA targeted cells, both in the BM and SP samples obtained from mice transplanted with SCID-X1 patient 1 and 2 derived CD34^+^ HSPCs **(Fig. 4e and Supplemental Fig. 12a)**. A statistically significant difference was observed only in BM samples derived from SCID-X1 patient 3 engraftment (\*\**P* = 0.0073, Holm-Sidak test) **(Supplemental Fig. 12d)**. Noteworthy is the observation that only the codon optimized *IL2RG* cDNA was detected in the SP of mice (n = 8) engrafted with SCID-Xl patient 2 corrected CD34^+^ HSPCs **(Table 2)**, consistent with the survival advantage that a cell with a corrected *IL2RG* gene has over a non-functional one. Multi-lineage analysis of SP samples derived from mice engrafted with *IL2RG* cDNA targeted SCID-Xl mPB CD34^+^ HSPCs derived from patient 2 showed that significant levels of erythroid, myeloid and lymphoid lineages were established **(Fig. 4f)** with the absolute numbers shown in **(Fig. 4g)**. We note that the lymphoid lineage was not established from *IL2RG* cDNA targeted SCID-X1 derived CD34^+^ HSPCs from patient 1 and 3 due to the overall low levels of human engraftment in mice transplanted with both mock and HR-GE modified cells **(Supplemental Fig. 12).** Engraftment and *in vivo* functional studies using frozen mobilized SCID-X1 patient derived and corrected CD34^+^ HSPCs are difficult to carry out. This is due to low overall human engraftment when using frozen mobilized CD34^+^ HSPCs and technical difficulties pertaining to intrahepatic delivery of the cells. As a result, insufficient number of T-cells were generated to run additional *in vivo* characterization. Despite this, our work is the first to show *in vivo* rescue of the lymphoid lineage in a SCID-X1 patient derived CD34^+^ HSPCs. In sum, these xenotransplantation studies demonstrated that *IL2RG* cDNA targeted HSPCs can engraft and rescue the SCID-X1 phenotype, as demonstrated by multi-lineage reconstitution both *in vitro* and *in vivo.* These results further highlight the effectiveness and benefit of a selection free genome targeting approach to treat SCID-Xl disease. Finally, we observed no abnormal hematopoiesis in mice transplanted with HR-GE patient derived cells providing further evidence for the safety of the process.

**Figure 4.**
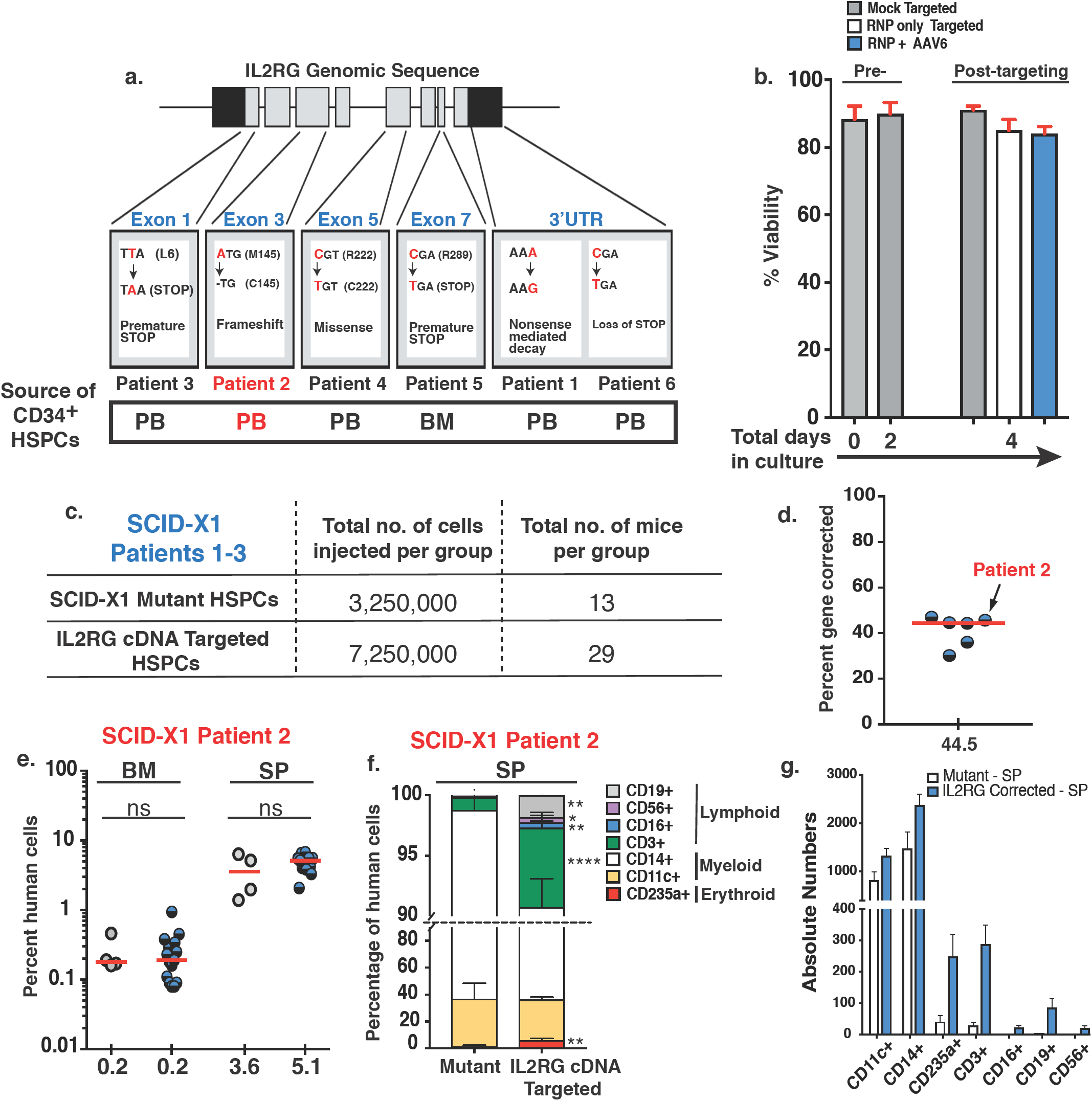
In vivo rescue of SCID-X1 mutation. **a.** Mapping and description of SCID-X1 mutations. **b.** Percent viability determined at indicated days pre- and post-targeting. Targeting conditions: mock (nucleofected only), RNP (nucleofected with RNP only), RNP + AAV6 (nucleofected with RNP and transduced with AAV6-based IL2RG corrective donor); Shown is data for mobilized peripheral blood CD34^+^ cells (n = 5). **c.** Summary of SCID-X1 derived CD34^+^ mutant and IL2RG cDNA targeted HSPCs transplanted into NSG pups. **d.** Day 2, medium-scale (1.0 × 10^6^) ex-vivo genome targeting frequencies of frozen mobilized peripheral blood SCID-X1 patient derived CD34^+^ HSPCs (blue-black circles) (n = 6). Arrow shows 45% genome targeting of SCID-X1 patient 2 derived CD34^+^ cells. **e.** Human cells engraftment analysis at week 17 after intra-hepatict delivery of IL2RG cDNA targeted (blue-black circles, n = 15) or mutant HSPCs (grey circles, n = 4). **f.** Percent cellular composition of the lymphoid, myeloid and erythroid lineage derived from IL2RG corrected or mutant HSPCs. CD3^+^: \*\*\*\**P*-value < 0.0001, CD56^+^: \**P*-value = 0.0146, CD16^+^: \*\**P*-value = 0.0013, CD19^+^: \*\**P*-value = 0.0015, CD235a^+^: \*\**P*-value = 0.0022 (Welch’s t-tes). RNP: ribonuclearprotein. **g.** Absolute numbers derived from **f.**

**Table 2.** Summary for primary human transplants of IL2RG cDNA targeted and IL2RG mutant SCID-X1 patient 2 derived CD34^+^ HSPCs.

| **Condition** | **Bone Marrow**<br>**(median with range)** | **Spleen**<br>**(median with range)** |
| --- | --- | --- |
| **Mutant HSPCs** | 0.18 (0.16 - 0.46) n = 4 | 3.6 (1.39 - 6.3) n = 4 |
| **IL2RG cDNA Targeted HSPCs** | 0.2 (0.08 - 0.94) n = 15 | 5.1 (2.1 - 6.7) n = 15 |
| **IL2RG cDNA Targeted in IL2RG**<br>**Bulk engrafted HSPCs** | QNS | 100.0 %<br>n = 8 |

### Signaling and proliferation of *IL2RG* cDNA targeted T-cells at endogenous start site

To address the receptor function and signaling in progenitor cells in which the gene is express through the targeted integration of a codon-optimized cDNA into the translational start site of the endogenous locus, we evaluated the proliferation and signaling activity of HR-GE human T lymphocytes derived from healthy male donors. Mature T cells depend on proper *IL2RG* expression and signaling through *IL2RG*-containing receptors, e.g., IL-2R, to promote proliferation and differentiation^39^. Activation of T-cells by CD3/CD28 antibodies leads to a rapid induction of IL-2 cytokine, which in turn signals though the IL-2R. Subsequent phosphorylation of tyrosine residues on the cytoplasmic domains of the receptors initiates a cascade of events that phosphorylate and activate the signaling transducers and activators of transcription 5 (STAT5) proteins. Therefore, we assessed the levels of pSTAT5 in *IL2RG* cDNA targeted T-cells, where the *IL2RG* cDNA donor contained *tNGFR* selectable marker. **(Fig. 5a, bottom flow chart)**. Intracellular staining for pSTAT5 from *IL2RG* cDNA targeted T-cells **(Fig. 5b, top panels)** and levels of pSTAT5 (ratio of tNGFR^+^pSTAT5^+^ double positive cells to that of tNGFR^+^ only cells, marked red) was demonstrated to be comparable to that of unmodified normal T-cells 69.3 ± 7.0 *vs* 67.7 ± 4.4 (mean ± *s.e.m*.), respectively **(Fig. 5c)**. As expected, knocking-out (KO) the *IL2RG* locus with an *IL2RG* targeted donor expressing only *tNGFR*, significantly reduced the levels of pSTAT5 (12.7 ± 5.6; mean ± SEM) **(Fig. 5b bottom panels and Fig. 5c)**.

**Figure 5.**
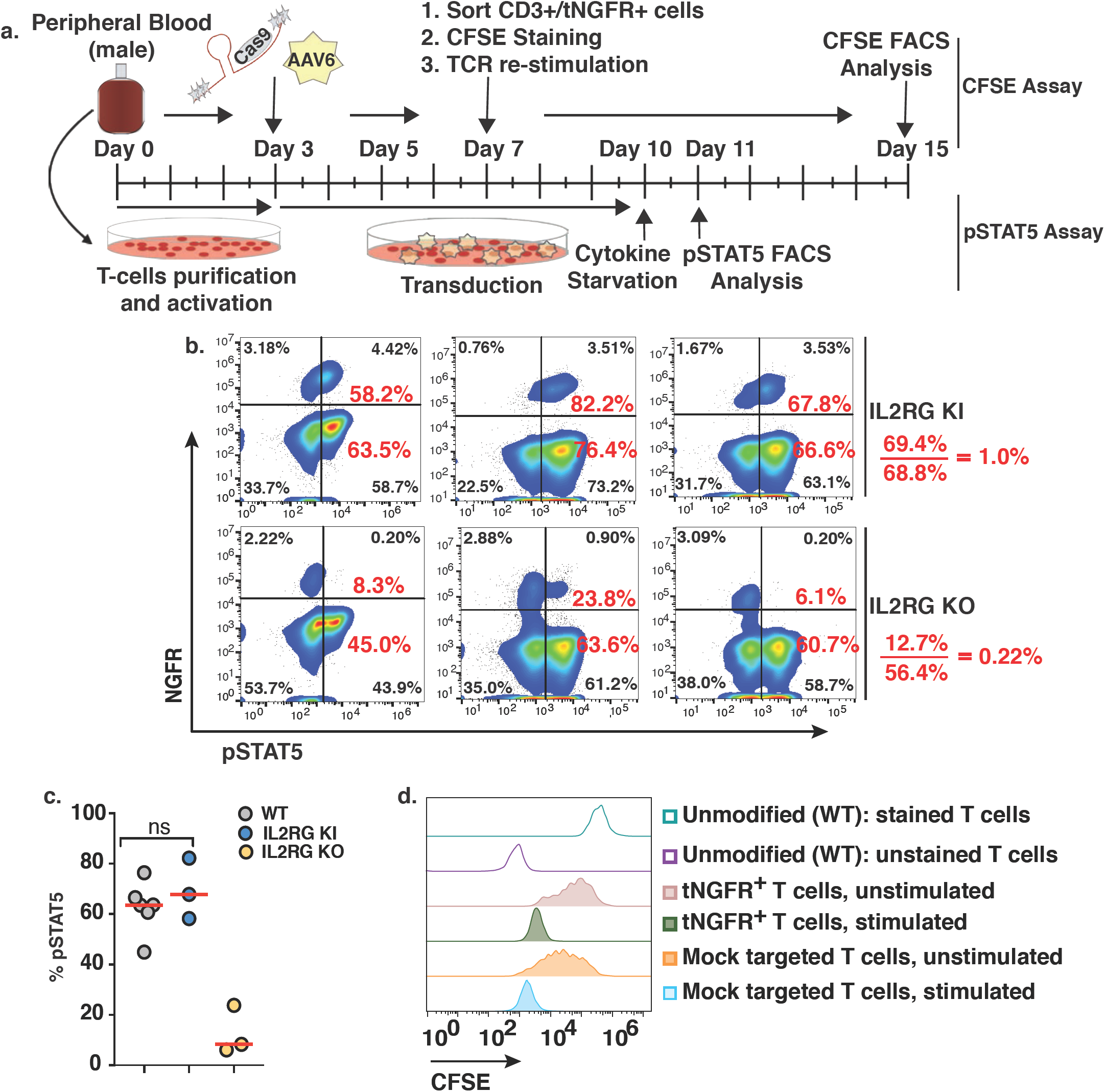
Evaluation of IL-2 receptor function in IL2RG cDNA targeted T-cells. **a.** Schematic of signaling (pSTAT5 - bottom) and proliferation (CFSE - top) in vitro assays. **b.** pSTAT5 assay derived FACS plots. Top: healthy male derived T-cells genome targeted with IL2RG cDNA tNGFR^+^ (KI) or with an tNGFR^+^ only cassette integrated at the IL2RG endogenous locus (KO). In red are the percent of double positive IL2RG-tNGFR^+^pSTAT5^+^ [4.42% / (4/42% + 3.18%)]*100. We compare 58.2% cells (IL2RG targeted T-cells) with 58.7% (IL2RG from WT T-cells). Shown are 3 biological replicates. **c.** Quantification of IL-2R signaling through STAT5 pathway. WT T-cells (grey circles, n=6), IL2RG KI (blue circles, n=3) and IL2RG KO (orange circles, n=3) **d.** Proliferation profile of CFSE labeled, TCR stimulated IL2RG cDNA tNGFR^+^ sorted or mock targeted T-cells. Mock targeted T-cells are WT T-cells cultured for the same amount of time as the NGFR^+^ targeted cells and have been nucleofected in the absence of RNP or absence of trasduction with AAV6. Shown FACS analysis at days 2, 4, 6 and 8. pSTAT5: phosphorylated STAT5; CFSE: carboxyfluorescein succinimidyl ester, KI: knocked in, KO: knocked out, tNGFR: truncated nerve growth factor receptor. IL-2: interleukin 2.

To demonstrate that the genome edited IL2R is permissive for proliferation upon engagement of IL-2 cytokine, we quantified the levels of proliferation of *IL2RG* cDNA targeted T-cell following T cell receptor (TCR) stimulation. A CFSE (carboxyfluorescein succinimidyl ester) dilution assay was used to measure whether targeted insertion of the codon-optimized cDNA could support T-cell proliferation. Loss of CFSE signal occurs when cells proliferate as the dye dilutes from cell division. An overview of the assay is shown **(Fig. 5a, top flow chart)**. In our experimental settings, we observed similar proliferation profile in *tNGFR*^+^ T-cells (marking cells in which the *IL2RG* cDNA had been knocked-in at the start codon) when compared to mock targeted cells **(Fig. 5d)**. We note that the “unmodified” cells had not undergone prior bead stimulation and so remained quiescent while the targeted and mock cells had undergone prior bead stimulation and thus there was residual proliferation without re-stimulation in those cells giving the broader peak. Overall, our data demonstrates that the genomic integration of an *IL2RG* codon diverged cDNA at the start site of the endogenous locus preserves the normal signaling and proliferation capacity of genome targeted human T-cells.

### Off-target and karyotype analysis of CRISPR/Cas9 genome editing platform

Gene therapy clinical trials for SCID-X1 demonstrated genotoxicity development due to insertional mutagenesis^6^. In the gene editing approach, the concern of random or semi-random integration is eliminated since integration of the cDNA is mediated through double-strand break mediated homologous recombination into a specific genomic target (in this case, exon 1 of the *IL2RG* gene). We investigated the fidelity of the dsDNA break generated by the CRISPR/Cas9 RNP complex, which could be a potential source of genotoxicity. The off-target activity of the full-length 20nt and three truncated versions (19nt, 18nt and 17nt) of sg-1 guide were assessed at 54 different potential sites predicted by either Guide-Seq in U20S cells^40^ or bioinformatically-COSMID^41^ **(Fig. 6a).** The analysis was performed in both healthy **(Fig. 6a)** and patient derived CD34^+^ HSPCs **(Fig. 6b)** to assess the specificity of the sg-1 gRNA. At the three sites identified by Guide-Seq analysis, there was no evidence of off-target INDELs. In the 51 sites identified by COSMID, only two showed evidence of off-target INDELs, both at levels <1% **(Fig. 6a)**. We detected INDEL frequencies using the 20nt sg-1 of 0.59%, in an intron of myelin protein zero-like 1(*MPZL1*), a cell surface receptor gene involved in signal transduction processes. The 19nt sg-1 induced a lower frequency of off-target INDELs 0.11% **(Fig. 6a and Table 3).** We also analyzed the INDEL frequencies of potential off-target sites in genome edited CD34^+^ HSPCs derived from SCID-Xl patient 1 in which the cells were edited using the 19nt sg-1 **(Fig. 6b)**. We found INDEL frequencies of 0.08% at *MPZL1* and 0.27% at the *ZFN330* site (intergenic and >9kb from the nearby gene, respectively) **(Table 3)**. Off-target activities of sg-1 guides, WT (20nt) and truncated (19nt), were further assessed in the context of a high fidelity (HiFi) Cas9 protein^42^ in SCID-X1 CD34^+^ HSPCs. While viability **(Fig. 6c)** and editing frequencies (% INDELs) **(Fig. 6d)** were at comparable frequencies between WT and HiFi Cas9, the *IL2RG* cDNA targeting frequency (%HR) was reduced slightly for both the 20nt IL2RG sg-1 guide (58% to 47.6%) **(Fig. 6e)** and the 19nt guide (44.3% to 39.6%) **(Fig. 6f)**. Using the 20nt sg-1 combined with the HiFi Cas9, however, resulted in no detectable INDELs (“background” Table 3) at the two sites for which there was low but detectable INDEL frequency (0.1% at MPZL1 and 0.27% at ZNF330) using the 19nt sg-1 combined with wild-type Cas9.

**Figure 6.**
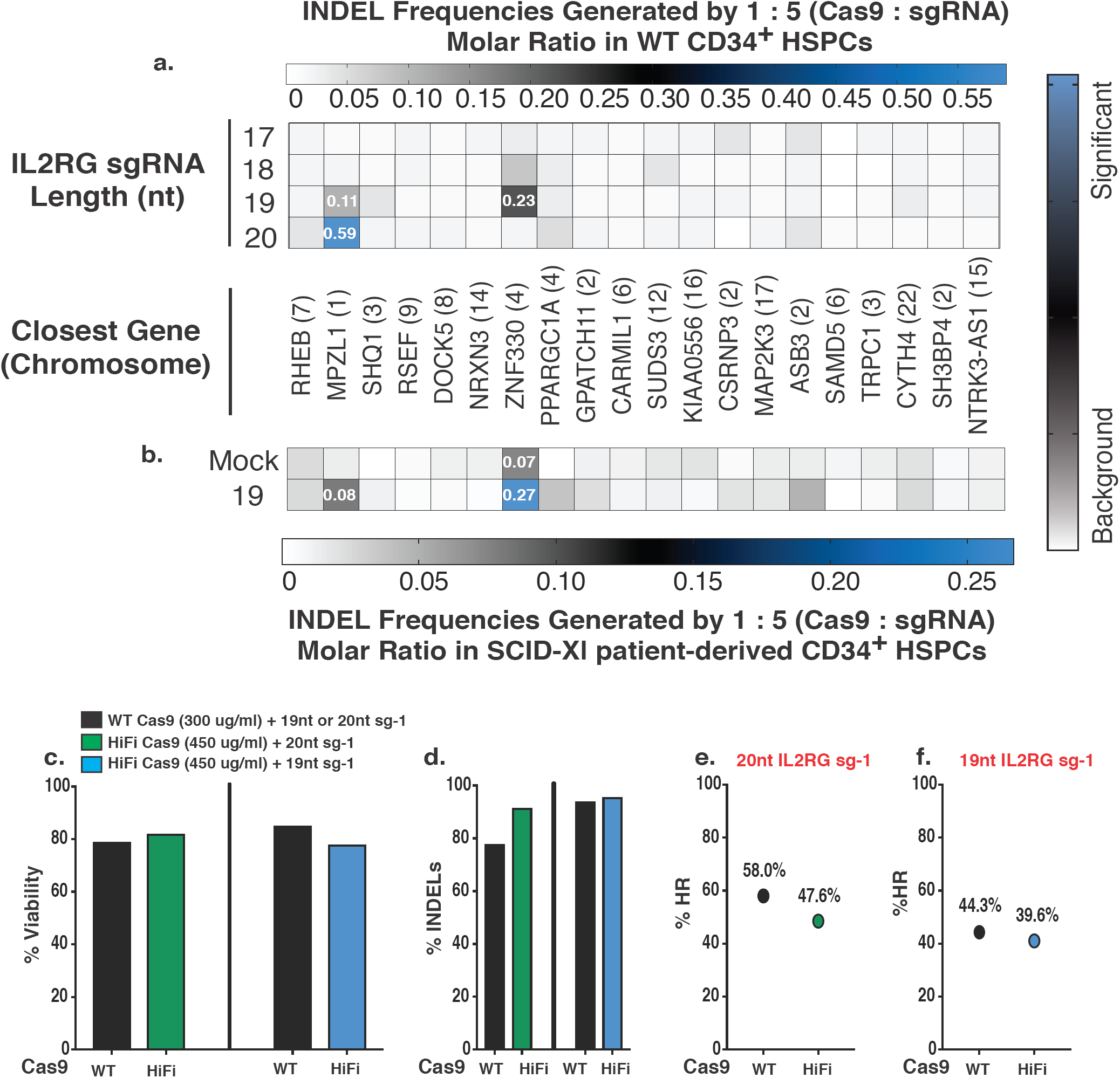
Genome specificity of IL2RG sgRNA guide. **a.** Heat map of off-target INDEL frequencies quantified by NexGen-Seq at COSMID identified putative off-target locations from healthy CD34^+^ HSPCs. Levels of NHEJ induced by 20nt IL2RG sgRNA and truncated 19nt, 18nt and 17nt pre-complexed with WT Cas9 protein at 5:1 molar ratio. **b.** Heat map as in (**a**) of off-target INDEL frequencies derived from 19nt IL2RG sgRNA #1 in the genome of CD34^+^ HSPCs SCID-X1 patient 1 derived cells. **c.** Percent viability at day 4 of SCID-X1 patient derived CD34^+^ HSPCs nucleofected with either wild type (WT) or high fidelity (HiFi) *Sp*Cas9 protein pre-complexed with either the 20nt or the 19nt IL2RG sg-1. **d.** Percent INDELs measured by TIDE analysis at day 4 using WT or HiFi Cas9 protein pre-complexed with the 20nt IL2RG sg-1(left panel) or 19nt IL2RG sg-1 (right panel). **e.** Percent IL2RG cDNA targeting (% HR) as measured by ddPCR at day 4 generated by either WT or HiFi Cas9 protein pre-complexed with the 20nt IL2RG sg-1 or **f.** 19nt IL2RG sg-1.

**Table 3.** Summary of IL2RG sgRNA off-target INDEL frequency analysis.

| Gene Name | Method of Identification |  | Chromosome Location | Feature | *Expression in Hematopoietic Stem Cells | U2OS | WT CD34<sup>+</sup> | SCID-X1 CD34<sup>+</sup> | SCID-X1 CD34<sup>+</sup> | SCID-X1 CD34<sup>+</sup> |
| --- | --- | --- | --- | --- | --- | --- | --- | --- | --- | --- |
|  | Guide Seq | COSMID |  |  |  | Plasmid | 20nt RNP (WT Cas9) | 19nt RNP (WT Cas9) | 20nt RNP (HiFi Cas9) | 19nt RNP (HiFi Cas9) |
| IL2RG |  |  | X | Exon | Yes | 81.1% | 81.7% | 91.7% | 94.1% | 97.6% |
| LIN01287 | ✓ |  | 7 | Intergenic | Data not available | background | background | background | n/s | n/s |
| MPZL1 |  | ✓ | 1 | Intron | Yes | 1.1% | 0.1% | 0.1% | background | background |
| SHQ1 | ✓ | ✓ | 3 | Intron | Yes | 1.5% | background | background | n/s | n/s |
| SMYD3 | ✓ |  | 1 | Intron | Yes | 4.2% | background | background | n/s | n/s |
| ZNF330 |  | ✓ | 4 | Intergenic | Data not available | background | 0.23% | 0.27% | background | background |

*\*Expression determined by www.biogps.org and Gene Expression Commons*

To further assess whether genomic instabilities, particularly translocations, were generated by the CRISPR/Cas9-AAV6 based genome editing process, we performed karyotype analysis on CB-derived CD34^+^ HSPCs from healthy male donors. We chose to run karyotype analysis over PCR-based translocation assays because we have previously found that the frequency of translocations in CD34^+^ HSPCs when two on-target breaks (with INDEL frequencies of >80%) was 0.2-0.5%^43^. The probability that there is a translocation between the on-target break and break that has an INDEL frequency of <0.1% is exceedingly low. Whole chromosomal analysis was performed on ≥ 20 cells from the different conditions (WT, mock only, RNP only, AAV6 only and RNP+AAV6). The analysis confirms absence of any chromosomal abnormalities in 20 out of 20 untreated or mock treated cells **(Supplemental Fig. 14a and 14b)**, 40 out of 40 RNP only or RNP treated with rAAV6 **(Supplemental Fig. 14c and 14d)** and from 40 out of 40 cells treated with rAAV6 only **(Supplemental Fig. 14d and 14e).** Lastly, we performed γH2AX and relative survival assays in K562 and 293T cells lines, respectively, to determine and compare the levels of DNA damage and toxicity induced by ZFN, TALEN and Cas9/gRNA nucleases that all target the *IL2RG* gene **(Supplemental Fig. 15a and 15b)**. The CCR5 ZFNs were first described in Perez *et al*^44^ (2008) and subsequently used clinically and to modify HSPCs^45,46,23^. The nucleases targeting the *IL2RG* gene were described previously in Urnov *et al*^25^ (2005) (ZFNs) and Hendel *et al*^*47*^ (2014) (ZFNs, TALENs, and CRISPR/Cas9). The Cas9/gRNA nuclease generated the lowest levels of toxicity by showing fewer γH2AX foci and higher percent survival of human cells over-expressing each nuclease^48^ highlighting the notion that standard TALEN and ZFN nuclease platforms are less specific than CRISPR/Cas9.

In conclusion, our off-target analysis confirms that high specificity and activity is achieved using the *IL2RG* CRISPR/Cas9-AAV6 HR-GE system described here.

## Discussion

Currently there are numerous genome-editing based clinical trials in USA and China, none of which are for the treatment of PIDs (clinicaltrials.gov). There have been a number of proof-of-concept genome editing studies to explore the feasibility and safety of using a homologous recombination (HR)-mediated approach to correcting pathologic mutations in the *IL2RG* gene as a path to developing an auto-HSPC based therapy for SCID-X1^*25, 18, 19, 49, 50*^. In particular, a recent study by Schiroli *et al*^*18*^, in the process of developing a clinical translation genome editing for SCID-Xl in CD34^+^ HSPCs, designed a ZFN GE-based platform to integrate a full *IL2RG* cDNA at intron 1 delivered by integration-defective lentiviral vector (IDLV) or rAAV6. They were able to generate ∼ 40% INDELs and ∼10% HDR frequencies in WT CD34^+^ HSPCs, with targeted integration frequencies of ∼25% in one SCID-Xl patient derived cord blood CD34^+^ HSPCs. In this work, Schiroli, J., *et al*^*18*^ performed only one experiment combining CRISPR/Cas9 with AAV6 and almost all of their data, including the engraftment data was from ZFN modified cells. Even in the engraftment studies, they did not demonstrate the engraftment potential of ZFN/AAV6 intron 1 *IL2RG* corrected SCID-X1 patient derived cells nor the *in vivo* rescue of multi-lineage developmental potential and lymphoid development from corrected patient derived cells as we have demonstrated in this work. Moreover, while the study in the mouse model clearly demonstrates that a LT-HSC correction frequency of ≥ 10% would be ideal, their NSG transplant studies using ZFN-IDLV edited cells did not reach this benchmark in either primary transplants where the median frequency of HR cells in HSPCs was ∼5% or in secondary transplants (median frequency of HSPC HR cells was ∼2%).

Here, we describe a CRISPR/Cas9-AAV6 genome targeting pre-clinical study where a full length *IL2RG* cDNA is precisely and efficiently integrated at the endogenous translational start site in CD34^+^ HSPCs. Compared to previous work by Dever, D.P. *et al*^*24*^, where the payload consisted of only a single nucleotide based corrective donor, our approach, despite being more challenging, delivers a median of 45% genome targeting with higher frequencies of 50-60% being achieved regularly. Our therapeutic approach has the benefit of not only being able to correct >97% of known SCID-X1 pathogenic mutations due to the “universal” strategy design used but also to achieve high levels, up to 20%, of *IL2RG* cDNA genome targeting frequencies in LT-HSCs, as demonstrated from secondary transplants studies. This high levels of genomic editing of LT-HSCs has not been reported, to date, in the literature and has great implication in advancing the next generation of genome editing based gene therapies. Therefore, out genome editing platform and methodology shows significant advancements over the ZFN genome editing approach or even current CRISPR/Cas9 (RNP)– AAV6 approach (Genovese, P. *et al*^19^, Dever, P.D., *et al*^*24*^, Schiroli J., *et al*^*18*^). This level of correction is likely to be curative based on both animal studies^18^, from patients who had spontaneous reversion mutations in progenitor cells and from human gene therapy clinical trials. In the gene therapy clinical trials for SCID-X1, immune reconstitution was achieved with as little as 1% of the cells having gene transfer^*3*^ or from vector copy numbers (VCN) of only 0.1 in the blood^2^. Our results also demonstrate a lack of functional toxicity from the CRISPR/Cas9-AAV6 procedure because LT-HSCs were preserved and because normal human hematopoiesis was obtained from the genome edited cells.

The safety of the approach is further supported by the lack of karyotypic abnormalities generated in RNP exposed CD34^+^ HSPCs and by INDEL frequencies below the limit of detection using a high-fidelity version of Cas9 at 54 potential off-target sites identified by bioinformatics and cell-based methods. Even using wild-type Cas9, off-target INDELs were only detected at two sites (both at low frequencies (<0.3%)) which were at sites of no known functional significance and did not result in any measurable perturbations in the cell population in all the assays used in this work the most important of which was no evidence of abnormal hematopoiesis in RNP treated cells. In the course of these studies we transplanted a full human dose for an infant (the target age that we are planning to treat in a phase I/II clinical trial) into NSG mice (28.4 million CD34^+^ HSPCs), a functional safety standard that the FDA has used prior to approving a phase I clinical trial of ZFN editing of CD34^+^ HSPCs^23^.

Our work lays the foundation pertaining not only to the technology and methodology required to establish a genome editing base gene therapy for SCID-X1 but it also presents the initial investigation towards its safety: off target activity of the sgRNA guide, hematopoietic engraftment potential and multi-lineage development. As a pre-clinical work, engraftment studies are the only way we can establish the initial stages of safety. At such, persistence of *IL2RG* genome targeted cells for 8 months in a surrogate system such as that of an immunodeficient mouse is the current gold standard, which we have achieved. While the ultimate test of the safety and efficacy of our approach will be established during a gene therapy clinical phase I/II trial, we believe that our conclusion of the safety of our proposed methodology, within the limits of a pre-clinical study, is valid.

In addition to showing the CRISPR/Cas9-AAV6 HR-GE system safely and effectively modifies healthy donor LT-HSCs, we also demonstrate that the approach can efficiently modify peripheral blood patient derived CD34^+^ HSPCs (residual cells released for research that were originally harvested as part of a lentiviral gene therapy trial) with a median correction frequency of 44.5%. We achieved this frequency without having to incorporate a selection strategy, which may have contributed to achieving higher frequencies of targeted integration than previously reported in LT-HSCs^19,24,18,51^. The correction of the patient derived cells rescued their lymphoid developmental potential using both *in vitro* and *in vivo* assays. Specifically, we have shown *in vitro* OP9-idll1 system **(Fig 1e**), *in vivo* engraftment studies using healthy donors CD34^+^ HSPCs **(Fig. 3a**, **Fig. 3b**, **Fig. S7)** and *in vivo* engraftment using SCID-X1 patient derived CD34^+^ HSPCs **(Fig. 4F)** that our genome editing approach using a full length *IL2RG* cDNA has succeeded in establishing the lymphoid lineage: T-cells, NK cells in addition to B-cells. Neither the work of Genovese, P. *et al*^*19*^ nor Schiroli, J., *et al*^*18*^ have demonstrated any level of human engraftment and lymphoid development from SCID-X1 patient derived cells. In addition, we have shown that our genome editing approach retains functionality and proper signaling when editing T-cells **(Fig.5b-d).** Moreover, the corrected patient derived cells gave equivalent engraftment following transplantation compared to unmanipulated cells demonstrating that the process had not significantly damaged the cells.

Unlike previous work by Dever, *et al*^*24*^ and Schiroli *et al*^*18*^ we chose to carry out xenotransplantation assays using un-enriched *IL2RG* genome targeting CD34^+^ HSPCs because of the selective growth advantage within the lymphopoietic compartment that a cell with a corrected *IL2RG* gene has over cells with a non-functional gene. In addition, we have demonstrated that higher frequencies of gene targeting occurs using the shorter donor vector without selection (a process that might inadvertently enrich for progenitor cells rather than stem cells)^19^. The removal of a selection step also simplifies the overall cell manufacturing process thus facilitating easier translation to clinical scale manufacturing needed to start a phase I/II clinical trial.

We highlight that our *IL2RG* gene targeted CD34^+^HSPCs involves a maximum of 4 days of *ex-vivo* culturing in addition to genome editing manipulation. Still, we obtained a robust multilineage development, 16 weeks post engraftment into NSG mice – a detailed example is shown **(Supplementary Fig. S7)**. Moreover, not only there is no statistical difference noted in the levels of CD3^+^ cells derived from WT, Mock or *IL2RG* cDNA targeted and engrafted CD34^+^ HSPC by the intra-femoral route **(Fig. 3a, IF, BM)**, but the levels of CD3^+^ obtained are higher than those previously published^52,53^.

To determine the percent of INDELs present in the *IL2RG* cDNA targeted and bulk engrafted cells, we genotyped the bone marrow, spleen and peripheral blood samples obtained from week 16 primary intra-femoral human engraftment. TIDE analysis of the un-targeted cells confirmed the presence of INDELs at a frequency of >90%. Further characterization of the INDEL spectrum demonstrates that high rates of frameshift mutations (+1, −11, and −13 INDEL spectrum) are being created in the majority of CD34^+^ HSPCs not targeted with *IL2RG* cDNA. Therefore, the results generated by the engraftment studies using CB derived CD34^+^ HPSCs from healthy male donors strongly suggests that the *IL2RG* cDNA targeted cells were the vast majority of cells driving the establishment of multi-lineages, further supporting the functionality and efficacy of our *IL2RG* codon optimize cDNA targeted at transcription start site.

Our rationale for developing a genome editing-based gene therapy for SCID-X1 is to provide a safe, efficient, precise and effective treatment option for patients. While it is encouraging that improved methods for allo-HSCT are being developed and that the early results using lentiviral based gene therapy for SCID has been shown to be safe and effective, the long-term safety, efficacy and limitations of these approaches remains to be determined. Thus, it is important to continue develop alternative strategies for curing patients with SCID-X1 using approaches that are less genotoxic (the mutational burden from genome editing is >100 fold less than for lentiviral based modification strategies by comparing the frequency of off-target INDELs to the frequency of uncontrolled lentiviral insertions). Ideally, multiple effective options will be available for patients, their families, and their treating physicians in the future thus giving them the opportunity to choose the approach that best fits their needs and circumstances. In sum, the safety and efficacy data presented in this study provides strong support for the clinical development of functional gene correction using the CRISPR/Cas9-rAAV6 genome editing methodology to establish a long lasting therapeutic, potentially curative, strategy beneficial to >97% of SCID-X1 patients.

## Author Contributions

M.P.D and M.H.P designed the experiments, M.P.D performed all the *in vitro* and *in vivo* experiments on both healthy and disease CD34^+^HSPCs, analyzed, interpreted data and wrote the manuscript together with M.H.P. V.A.W performed all the *in vitro* T-cell experiments, data analysis and interpretations. B.T.D. assisted with OP9-idll1 and human engraftment experiments and analysis. W.S and C.S assisted with all human engraftment experiments. S.M purified CD34^+^HSPCs from healthy cord blood, assisted with primary and secondary experiments and purified CD34^+^HSPCs from SCID-Xl patient 4. C.E.N performed all the intra-femoral engraftments and obtained all the intra-femoral aspirates and peripheral blood samples from week 8 and 12 post-engraftment analysis. C.L and G.B. performed the off-target analysis on healthy and disease samples. E.J.K performed the γ-H2AX and survival assay in K562. N.P. performed a pilot screen for *IL2RG* sgRNA guides and identified sgRNA guide 1. M.A.I developed the OP9-idll1 system and N.S. further optimized it for human CD34^+^ HSPCs studies, under the K.I.W. guidance. K.I.W provided the bone marrow derived SCID-X1 patient sample and thoughtful discussions support and advice on the manuscript. S.S.D and H.M. provided the frozen mobilized peripheral blood SCID-X1 patient samples and assisted in analyzing the the data from editing patient derived cells. M.G.R provided helpful discussions and input into data analysis and assessment of functional rescue of using the cDNA knock-in strategy.

## Acknowledgments

V.A.W is supported by fellowships from the Care for Rare Foundation (Munich) and Deutsche Forschungsgemeinschaft (DFG). M.H.P gratefully acknowledges the support from CIRM (PC1-08111), the NIH/NIAID (R01 AI097320-01), the Laurie Kraus Lacob Translational Research Fund, the Amon Carter Foundation, and the Sutardja Family Fund for supporting this work. We thank Mr. Dana Bangs from Stanford Cytology Labs for karyotype analysis. We gratefully acknowledge Dr. Dullei Min and Damoun Torabi for their assistance with xenotransplantation studies. We thank Dr. Robertson Parkman (Stanford University), Dr. Annalisa Lattanzi and the entire Porteus Laboratory for useful discussions and support.

## Material and Methods

### CRISPR-Cas9 sgRNA

Seven *IL2RG*, exon 1 specific, 20 nucleotide length oligomer sequences, used in the initial screen, were identified using the online CRISPOR software (crispor.terof.net) and synthesized (Synthego, Redwood City, CA, USA) as part of a chimeric 100 nucleotides single guide RNA (sgRNA). Chemically modified sgRNA oligomers were manufactured using a proprietary synthesizer by Synthego Corp. (Redwood City, CA, USA) on controlled-pore glass (AM Chemicals, Carlsbad, CA, USA) using 2’-O-t-butyldimethylsilyl-protected and 2’-O-methyl ribonucleotide amidites (ChemGenes, Wilmington, MA) according to established procedures. Standard ancillary reagents for oxidation, capping and detritylation were used (EMD Millipore, Cincinatti, OH). Formation of internucleotide phosphorothioate linkages was performed using ((dimethylaminomethylidene) amino-3H-1,2,4-dithiazoline-3-thione (DDTT, ChemGenes, Wilmington, MA).

A set of 2’-*O*-methyl 3’phosphorothioate MS (*23*) modified full-length 20 nucleotides and three additional versions having 1, 2 and 3 nucleotides removed from the 5’ end of the complementary region of the *IL2RG* sgRNA guide #1 were synthesized (TriLink Biotechnologies, San Diego, CA, USA) and purified using reverse phase high-high performance liquid chromatography (HPLC). Purity analysis was confirmed by liquid chromatography – mass spectrometry (LC-MS).

**Table S1:**
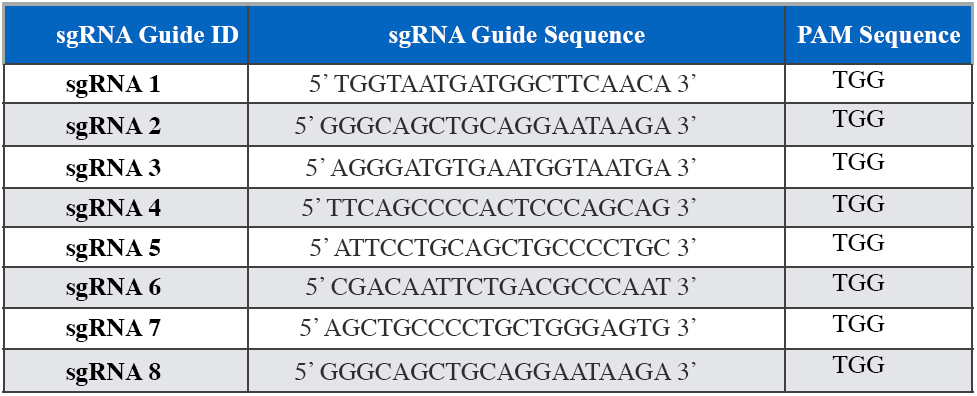
sgRNA guides for *IL2RG* exon 1.

### AAV6-based DNA donor design and vector production

All homology based AAV6 vector plasmids were cloned into pAAV-MCS plasmid containing AAV2 specific ITRs (Stratagene now part of Agilent Technologies, Santa Clara, CA, USA) using Gibson Assembly cloning kit according to the instructions in the commercial kit (New England Biolabs, cat # E5510S). Corrective, codon diverged *IL2RG* cDNA was designed to contain silent mutations that generated 78% sequence homology to the endogenous, wild type gene. All AAV6 viruses were produced in 293T in the presence of 1ng/ml sodium butyrate (Sigma-Aldrich, cat. no. 303410) cells and purified 48h later using an iodixanol gradient approach as previously described (*5*).

### CD34^+^ hematopoietic stem and progenitor cells

Mobilized peripheral blood and bone marrow CD34^+^ HSPCs cells were purchased from AllCells (Alameda, CA, USA). Cells were thawed using published protocol (^*54*^). Freshly purified cord blood-derived CD34^+^ HSPCs, of male origin, were obtained through the Binns Program for Cord Blood Research at Stanford University, under informed consent. Mononuclear cells isolation was carried out by density gradient centrifugation using Ficoll Paque Plus (400xg for 30 minutes without brake). Following two platelet washes (200 x g, 10-15 minutes with brake) HSPCs were labeled and positively selected using the CD34^+^ Microbead Kit Ultrapure (Miltenyi Biotec, San Diego, CA, USA) according to manufacturer’s protocol. Enriched cells were stained with APC anti-human CD34 (Clone 561; Biolegend, San Jose, CA, USA) and sample purity was assessed on an Accuri C6 flow cytometer (BD Biosciences, San Jose, CA, USA). Following purification or thawing, CD34^+^ HSCPs were cultured for 36-48 hours at 37°C, 5% CO_2_ and 5% O_2_, at a density of 2.5 × 10^5^ cells/ml in StemSpan SFEM II (Stemcell Technologies, Vancouver, Canada) supplemented with SCF (100 ng/ml), TPO (100ng/ml), Flt3-Ligand (100ng/ml), IL-6 (100 ng/ml), StemRegenin 1 (SR1) (0.75 mM) and UM171 (35 nM, Stemcell Technologies).

For secondary engraftment studies, CD34^+^ HSPCs were purified from total bone marrow of NSG mice at end point analysis. Sufficiently pure samples (greater than or equal to 80% CD34^+^) were pooled and cultured at 37°C, 5% CO_2_, and 5% O_2_ for 12 hours prior to secondary transplant.

### T-cells purification

Primary human T cells were obtained from healthy male donors from Stanford University School of Medicine Blood Center and purified by Ficoll density gradient centrifugation followed by red blood cell lysis in ammonium chloride solution (Stemcell Technologies, Vancouver, Canada) and magnetic negative selection using a Pan T cell isolation kit (Miltenyi Biotec, San Diego, CA, USA) according to manufacturer’s instructions. Cells were cultured at 37°C, 20% O_2_ and 5% CO_2_ in X-Vivo 15 (Lonza, Walkersville, MD, USA) supplemented with 5% human serum (Sigma-Aldrich, St. Louis, MO, USA) and 100 IU/ml human recombinant IL-2 (Peprotech, Rocky Hill, NJ, USA) and 10 ng/ml human recombinant IL-7 (BD Biosciences, San Jose, CA, USA). Cells were stimulated with immobilized anti-CD3 (OKT3, Tonbo Biosciences, San Diego, CA, USA) and with soluble anti-CD28 (CD28.2, Tonbo Biosciences) for three days prior to electroporation.

### Genome editing and INDEL quantification

Editing of all primary cells was carried out using a ribonucleic protein (RNP) system at a molar ratio of either 1:2.5 or 1:5 (Cas9 : sgRNA), unless otherwise stated. Recombinant S. *pyogenes* Cas9 protein was purchased from IDT (Integrated DNA Technologies, Coralville, Iowa, USA). Nucleofection was performed in P3 nucleofection solution (Lonza) and Lonza Nucleofector 4d (program DZ-100). Cells were plated at a concentration of 1.0×10^5^ – 2.5×10^5^ cells/ml. For T-cells editing, electroporation was performed using Lonza Nucleofector 4d (program EO-115) with an RNP composition as used for CD34^+^ HSPCs editing. INDEL frequencies were quantified using TIDE online software on genomic DNA extracted using Quick Extract (Epicentre, an Illumina Company, cat no. QE09050) according to manufacturing specifications.

### Genome targeting and quantification

CD34^+^ HSPCs nucleofected with the *IL2RG* specific RNP system were plated at a density of 5.0 × 10^5^ cells/ml and transduced with the AAV6 donor at an MOI of 200,000 vg/ul within 15 minutes of nucleofection. Cells were cultured at 37°C, 5% CO_2_, 5% O_2_ for 36h to 48h after which they were either re-plated in fresh media, at a density of 2.5 × 10^5^ cells/ml or prepared for xenotransplantation studies.

Absolute quantification of the levels of genomic integration was carried out using Digital Droplet PCR^™^ (ddPCR^™^, BioRad, Hercules, CA, USA). Genomic DNA was extracted as described in previous section. 1ug of genomic DNA was digested with EcoRV-HF (20U) in Cutsmart buffer at 37°C for 1h. ddPCR reaction contains 1x reference primer/probe mix synthesized at a 3.6 ratio (900 nM primer and 250 nM FAM labeled probe), 1x target primer/probe mix synthesized at a 3.6 ratio (HEX labeled probe), 1x ddPCR Supermix for probe without dUTP, 50 ng of digested DNA and water for a total volume of 25 ul. The primers and probes sequences are detailed in Table S2.

Genomic DNA in the ddPCR mixture was partitioned into individual droplets using QX100 Droplet Generator, transferred to a 96 deep well PCR plate and amplified in a BioRad PCR thermocycler. The following ddPCR program was optimized to amplify a 500 bp amplicon: step1 - 95°C for 10 min, ramp 1°C/sec, step2 - 94°C for 30 sec, ramp 1°C/sec, step3 - 60.8°C for 30 sec, ramp 1°C/sec, step4 - 72°C for 2 min, ramp 1°C/sec, step5 - repeat step 2-4 for 50 cycles, step6 – 98°C for 10 min, ramp 1°C/sec, step7 – 4°C, ramp 1°C/sec. BioRad Droplet Reader and QuantaSoft Software were used to read and analyzed the experiment following manufacturer’s guidelines (BioRad). Absolute quantification as copy of DNA/ul was determined for the reference, endogenous *IL2RG* gene and for the integrated *IL2RG* cDNA. Percent targeting in total population was calculated as a ratio of HEX to FAM signal. For all targeting experiments, genomic DNA was derived from male donors.

Quantification of *IL2RG* cDNA targeted integration frequencies in SCID-Xl patients was assessed based on agarose gel quantification as *IL2RG* cDNA signal ratio intensity.

**Table S2.**
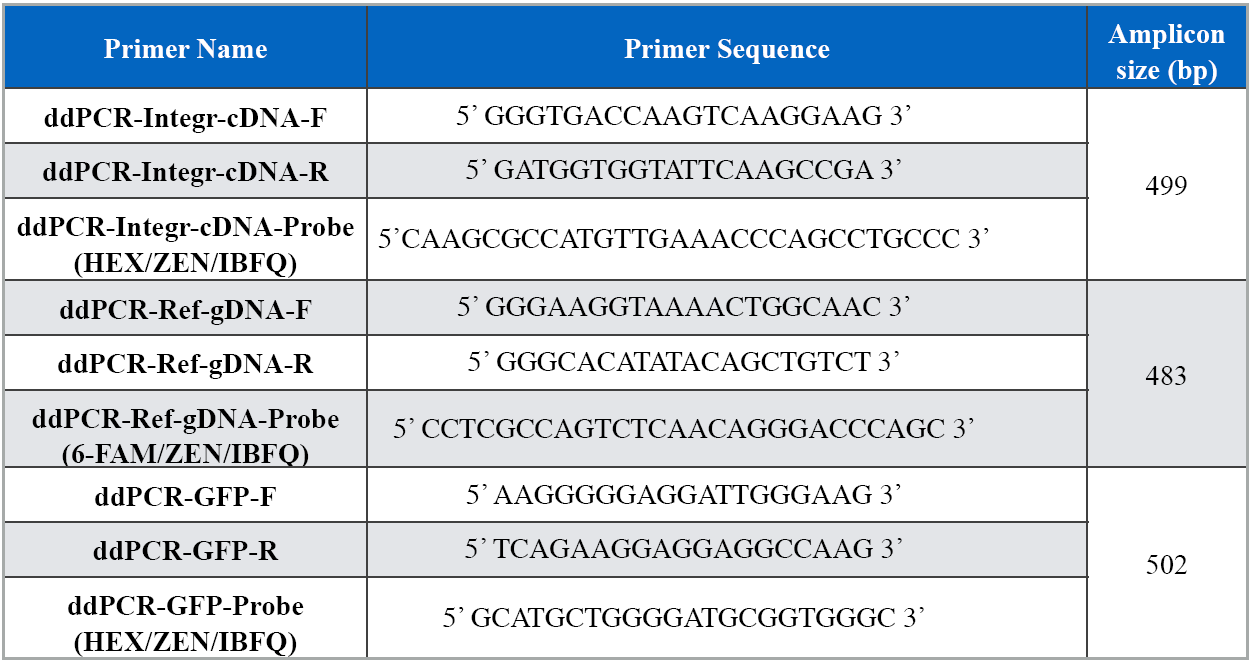
Primer information for ddPCR based assay.

### Methylcellulose colony-forming unit (CFU) assay

Two days post genome targeting single cells were sorted onto 96-well plates coated with MethoCult Optimum (StemCell Technologies, cat no H4034). 14 days later, colonies derived from targeted and mock treated cells were counted and scored based on morphological features pertaining to CFU-E, BFU-E, CFU-GM and CFU-GEMM. Genotyping analysis was performed to quantify the percent of mono-allelic targeting. A three primer-based *IL2RG* specific genotyping PCR based protocol was established an optimized as follows: *IL2RG* WT-F1 5’ 5’ GGGTGACCAAGTCAAGGAAG 3’; int-*IL2RG*-R1: 5’ GATGGTGGTATTCAAGCCGACCCCGA 3’; *IL2RG* WT-R2: 5’ AATGTCCCACAGTATCCCTGG. The PCR reaction contained 0.5 uM of each of the three primer, 1x Phusion Master Mix High Fidelity, 150ng - 200ng of genomic DNA and water to a final volume of 25 ul. The following PCR program generated an integration band of 543 bp from F1 and R1 primer set and an endogenous band of 1502 bp from F1 and R2 primer set: step 1 - 98°C for 30sec, step 2 - 98°C for 10sec; step 3 - 66°C for 30sec; step 4 - 72°C for 30sec, step 5 - repeat steps 2-4 for a total of 30 cycles, step 6 - 72°C for 7 min; step 7 - 4°C.

### OP9-idll1 system

ODT cells were generously provided by Dr. Irving Weissman’s lab and generated as previously described (*39*). Briefly, OP9 stromal cells were infected with two lentiviral constructs, the first containing a TET-ON tetracycline trans-activator (rtTA3) under control of a constitutive promoter (EF1a) and linked to turboRFP, and the second containing the Dll1 gene under control of a tet-responsive element (TRE) promoter and linked to turboRFP. In the presence of tetracycline or doxycyline, the rtTA3 rapidly activates expression of Dll1 and turboRFP.

### Lymphoid differentiation of SCID-X1 patient derived CD34^+^ HSPCs using OP9-idll1 *in vitro* assay

SCID-X1 patient derived CD34^+^ HSPCs were targeted with the *IL2RG* cDNA corrective donor. 48h post-targeting, 300 cells derived from either un-target or *IL2RG* cDNA targeted were sorted onto a well of a 96 well plate seeded with 50,000 OP9-idll1 cells 48h in advance. Cells were incubated at 37°C, 5% CO_2_, 10% O_2_ for one week in activation media containing: alpha-MEM base media (ThermoFisher, cat no. 32561102), supplied with 10% FBS (GemCell, cat no. 100-500), mono-thioglycerol (MTG) (100 uM), Ascorbic acid (50 ug/ml), 1x Penicillin/Streptomycin, SCF (10 ng/ml, PeproTech, cat no. AF-300-07), Flt-3L (5ng/ml, PeproTech, cat no. AF-300-19), IL-7 (5ng/ml PeproTech, cat no. 200-07), IL-3 (3.3 ng/ml, PeproTech cat no. AF-200-03), GM-CSF (10ng/ml, PeproTech, cat no. AF-300-03), TPO (10ng/ml, PeproTech cat no. AF-300-18), EPO (2U/ml, PeproTech, cat no. 100-64), IL-15 (10ng/ml, PeproTech cat no. AF-200-15), IL-6 (10ng/ml, PeproTech, cat no. 200-06). After 7 days, half the media was exchanged and doxycycline was added at a final concentration of 1ug/ml.

### *In vitro* multi-lineage differentiation analysis

Lymphoid, myeloid and erythroid differentiation potential was determined using FACS analysis at one week post dox induction. 100% growth was obtained from all wells seeded with 300 targeted or mock treated cells. Media was removed from all positive wells and cells were washed in 1X PBS. Cells were re-suspended in 50 ul MACS buffer (1x PBS, 2% FBS, 2mM EDTA), blocked for non-specific binding (5% vol/vol human FcR blocking reagent, Miltenyi cat no. 130-059-901), stained for live dead discrimination using Live/Dead blue dead cell staining kit for UV (ThermoFisher Scientific, cat no. L23105) and stained (30 min, 4°C dark) using CD3 PerCP/Cy5.5 (HiT3A, BioLegend), CD4 BV650 (OKT4, BioLegend), CD8 APC (HiT8a, BioLegend), CD11c BV605 (3.9, BioLegend), CD14 BV510 (M5E2, BioLegend), CD19 FITC (HIB19, BioLegend), CD33 AF-300 (WM53, BDPharmingen), CD45 BV786 (BDPharminge), CD56 PE (MEM-188 BioLegend), CD235a PE-Cy7 (HI264, BioLegend), CD271 (tNGFR) CF-594 (C40-1457, BD Horizon).

### Phosphorylated STAT5 *in vitro* assay on *IL2RG* targeted human T-cells

To assess STAT5 phosphorylation in response to cytokine stimulation, purified human T-cells were cultured for 7 days post electroporation and starved, overnight, in medium lacking serum and cytokines. Samples were split and either stimulated with IL-2 (100U/ml) and IL-7 (10ng/ml) or left un-stimulated. Cells were split again, fixed, permeabilized using 4% PFA and methanol and stained with CD3 PE (UCHT1, BioLegend), CD271 (tNGFR) APC (ME20.4, Biolegend). Intracellular antigens were stained with pSTAT5 AF-488 (pY694, BD Bioscience) or isotype control (BD Biosciences). FACS analysis was performed on Accuri C6 (BD Biosciences) or Cytoflex (Beckman Coulter) and data analysis was performed using FlowJo.

### CFSE cellular proliferation of *IL2RG* targeted human T-cells

Purified human T-cells were nucleofected alone (mock treated) or in the presence of the long corrective *IL2RG* cDNA-tNGFR DNA donor vector. NGFR^bright^ T-cells were sorted. NGFR^bright^ or mock treated cells were labeled with CFSE (BioLegend) according to the manufacturer’s protocol and either re-stimulated with Anti-CD3/Anti-CD28/IL-2/IL-7 as described in previous section or left unstimulated (IL-7 only). Targeting levels were monitored and quantified based on the tNGFR expression and on absolute quantification of the integrated *IL2RG* cDNA by ddPCR.

### Xenotransplantation of genome targeted CD34^+^ HSPCs into NSG mice

For all human engraftment studies we used freshly purified cord blood (CB) derived CD34^+^ HSPCs derived from healthy male donors, under informed consent. Human engraftment studies designed to rescue the disease phenotype were carried out using frozen, mobilized peripheral blood (mPB) CD34^+^ HSPCs derived from SCID-Xl patients 1-3. SCID-X1 patients were given subcutaneous injections of G-CSF (filgrastim, Neupogen®; Amgen, Thousand Oaks, CA) for 5 consecutive days at 10-16 mcg/kg/day and one dose of Pleraxifor for mobilization and apheresis (National Institutes of Allergy and Infectious Disease IRB-approved protocol 94-I-0073). Peripheral blood CD34^+^ HSPCs were selected from the leukepheresis product using Miltenyi CliniMACS.

Human engraftment experimental design and mouse handling followed an approved Stanford University Administrative Panel on Lab Animal Care (APLAC). Cells used for engraftment studies were exposed to a maximum of 4 days *ex-vivo* culturing.

### Intra-hepatic primary (1°) human engraftment

1.0×10^5^ to 2.5×10^5^ cells derived from *IL2RG* cDNA targeted cells or mock treated cells (electroporated in the absence of RNP and never exposed to AAV6) were re-suspended in 25ul - 30ul of freshly prepared CD34^+^ complete media with the addition of UM171 and SR1.

3-4 days old NSG pups were irradiated with 100 cGy and immediately engrafted intra-hepatic (IH) using an insulin syringe with a 27 gauge x 1/2” needle. A total of 2.15 × 10^6^ cells from each condition were injected into 11 pups/condition. 18/22 engrafted pups were analyzed at week 16 post-engraftment.

Level of human engraftment was assessed at weeks 8 and 12 using bone marrow aspirates and peripheral blood samples. At week 16 or later end point analysis was done from total bone marrow, spleen, liver and peripheral blood. For total bone marrow analysis, mouse bones were harvest from tibiae, femurs, sternum and spinal cord from each mouse and grinded using a mortar and pestle. Mononuclear cells (MNC) were purified using Ficoll gradient centrifugation (Ficoll-Paque Plus, GE Healthcare, Sunnyvale, CA, USA) for 25 min at 2,000 x g, at room temperature. Spleen and liver samples were grinded against a 40uM mesh, transferred to a FACS tube and spun down at 300 x g for 5 min, at 4°C. Red blood cells were lysed following a 10-12 min incubation on ice with 500ul of 1x ACK lysis buffer (ThermoScientific, cat no. A1049201). Reaction was quenched and cells were washed with MACS buffer (2% - 5% FBS, 2mM EDTA and 1x PBS). Peripheral blood samples were treated with 500 ul of 2% Dextren and incubated at 37C for 30 min to 1h. 800 ul to 1ml of the top layer was transferred to a FACS tube, spun down at 300 x g, 5 min and red blood cells lysed as already described.

Cells purified from all 4 sources were re-suspended in 50 ul MACS buffer, blocked, stained with LIVE/Dead staining solution and stained for 30 min at 4°C, dark with the following antibody panel: CD3 PerCP/Cy5.5 (HiT3A, BioLegend), CD19 FITC (HIB19, BioLegend), mCD45.1 PE-Cy7 (A20, BioLegend), CD16 PE-Cy5 (3G8, BDPharmingen), CD235a PE (HI264, BioLegend), HLA A-B-C APC-Cy7 (W6/32, BioLegend), CD33 AF-300 (WM53, BDPharmingen), CD8 APC (HiT8a, BioLegend), CD45 BV786 (HI3a, BD Horizon), CD4 BV650 (OKT4, BioLegend), CD11c BV605 (BioLegend), CD14 BV510 (M5E2, BioLegend) and CD56 Pacific Blue (MEM-188, BioLegend).

### Intra-femoral primary (1°) human engraftment

5.0×10^5^ cells derived from WT cells, mock treated, RNP treated and *IL2RG* cDNA targeted cells were injected intra-femoral (IF) into 6-8 weeks old NSG mice. Mice were irradiated with 200 cGy 2-4 hours prior to engraftment. Cells were prepared in the same fashion as described in the IH section. A total of 2.0×10^6^ WT cells were injected into a total of 4 mice, 3.5×10^6^ mock treated cells were injected into 7 mice, 2.0×10^6^ RNP treated cells were injected into 4 mice and 7.5×10^6^ *IL2RG* cDNA targeted cells were injected into 15 mice. 29/30 injected mice were analyzed at week 16 post-engraftment, as described in the IH engraftment assay section.

### Secondary (2°) human engraftment

Secondary engraftments experiments were derived from both IH and from IF engrafted human cells. From the IH mock and *IL2RG* cDNA targeted engrafted mice, total bone marrow was collected at week 16 post-primary engraftment, MNC were purified using Ficoll gradient centrifugation and CD34^+^ cells were enriched using CD34^+^ microbeads (Miltenyi). Enriched cells were pooled from 5 mock treated cells and from 7 *IL2RG* cDNA targeted cells and cultured overnight in complete CD34^+^ media containing UM171 and SR1. Following overnight incubation, cellular count and viability was determined for mock treated cells to be 2.47× 10^6^ cells at 85.5% viability and for *IL2RG* cDNA targeted cells was 4.8×10^6^ cells at 84% viability. 3.5×10^5^ mock treated cells and 5.0×10^5^ *IL2RG* cDNA targeted cells were engrafted IF into 8 6-8 weeks old, irradiated NSG mice (4 males and 4 females).

Secondary engraftment experiments derived from IF primary engraftments were carried on as described above with the following modification: 5.0×10^5^ CD34^+^ enriched cells derived from WT, Mock and RNP primary engraftment assay were IF injected into 4 6-8 weeks old NSG mice, 5.0×10^5^ CD34^+^ enriched cells derived from *IL2RG* cDNA targeted cells were IF injected into 12 6-8 weeks old NSG mice. Equal numbers of male and female mice were used.

### Intra-hepatic primary (1°) human engraftment of SCID-Xl patient CD34^+^ HSPCs

Frozen mobilized peripheral blood CD34^+^ HSPCs derived from SCID-Xl patients were thawed and genome targeted as described in previous section. 2.5×10^5^ cells were IH injected into 3-4 days old, irradiated NSG pups.

### GUIDE-Seq

sgRNAs were generated by cloning annealed oligos containing the IL2RG target sequence into pX330 (Gift from Feng Zhang, Addgene #42230)^*55*^. 200,000 U2OS cells (ATCC #HTB-96) were nucleofected with 1ug of pX330 Cas9 and gRNA plasmid and 100 pmol dsODN using SE cell line nucleofection solution and the CA-138 program on a Lonza 4D-nucleofector. The nucleofected cells were seeded in 500uL of McCoy’s 5a Medium Modified (ATCC) in a 24-well plate. Genomic DNA (gDNA) was extracted three days post-nucleofection using a Quick-DNA Miniprep plus kit (Zymo Research). Successful integration of the dsODN was confirmed by RFLP assay with NdeI. 400 ng of gDNA was sheared using a Covaris LE220 Ultrasonicator to an average length of 500bp. The sheared DNA was processed as previously described^*40*^ and sequenced on the Illumina Miseq. We analyzed GUIDE-Seq data using the standard pipeline^*40*^ with a reduced gap penalty for better detection of off-target sites containing DNA or RNA bulges.

### Bioinformatic off-target identification

Potential off-target sites for the IL2RG gRNA in the human genome (hg19) were identified using the web tool COSMID ^*41*^ with up to 3 mismatches allowed in the 19 PAM proximal bases. After off-target site ranking, 45 sites were selected for off-target screening.

### Off-target validation

Frozen mobilized peripheral blood CD34^+^ cells (AllCells) were electroporated with 300 ug/ml of Cas9 and 160 ug/ml of sgRNA. sgDNA was extracted 48h after RNP delivery. Off-target sites were amplified by locus-specific PCR. PCR primers contained adapter sequences to facilitate amplicon barcoding via a second round of PCR as previously described ^*56*^. All amplicons were pooled at an equimolar ratio and sequenced on the Illumina Miseq according to manufacturer’s instructions using custom sequencing primers for Read 2 and Read Index. Sequencing data was analyzed using a custom INDEL quantification pipeline ^*57*^.

### Karyotype analysis of *IL2RG* genome edited and cDNA targeted CD34^+^ HSPCs

Fresh cord blood CD34^+^ HSPCs were purified, genome edited or targeted as previously described. Four days post *ex-vivo* culturing and manipulations, 5×10^5^ cells from WT untreated, mock, RNP only, RNP and AAV6 or AAV6 only treated cells were processed by Stanford Cytology Labs at Stanford University. Karyotyping analysis was performed on 20 cells derived from each condition.

### IL2RG specific genotoxicity assays in human cell lines

Levels of γH2AX induced by different classes of engineered nucleases were quantified by measuring the phosphorylation of histone H2AX, a marker of DSB formation. K562 cells were nucleofected with the indicated doses of each nuclease expression plasmid, and the percentage of γH2AX^+^ cells was measured by FACS at 48h post-nucleofection. 293T cells were co-transfected with plasmids expressing GFP and nuclease. GFP positive cells were at day 2 and again at day 6 by FACS. Percent survival relative to I-SceI control was calculated as follows:

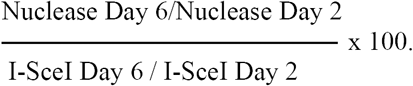

A percent equal to 100 denotes no toxicity while a percentage less than 100 marks toxicity.

### FACS analysis

All FACS analysis pertaining to OP9-idll1 and human engraftment analysis were done on FACS Aria II SORT instrument part of FACS Facility Core from Stanford University, Institute for Stem Cell Biology and Regenerative Medicine.

### Statistical analysis

Statistical analysis was done with Prism 7 (GraphPad Software).

---

**Fig. S1.**
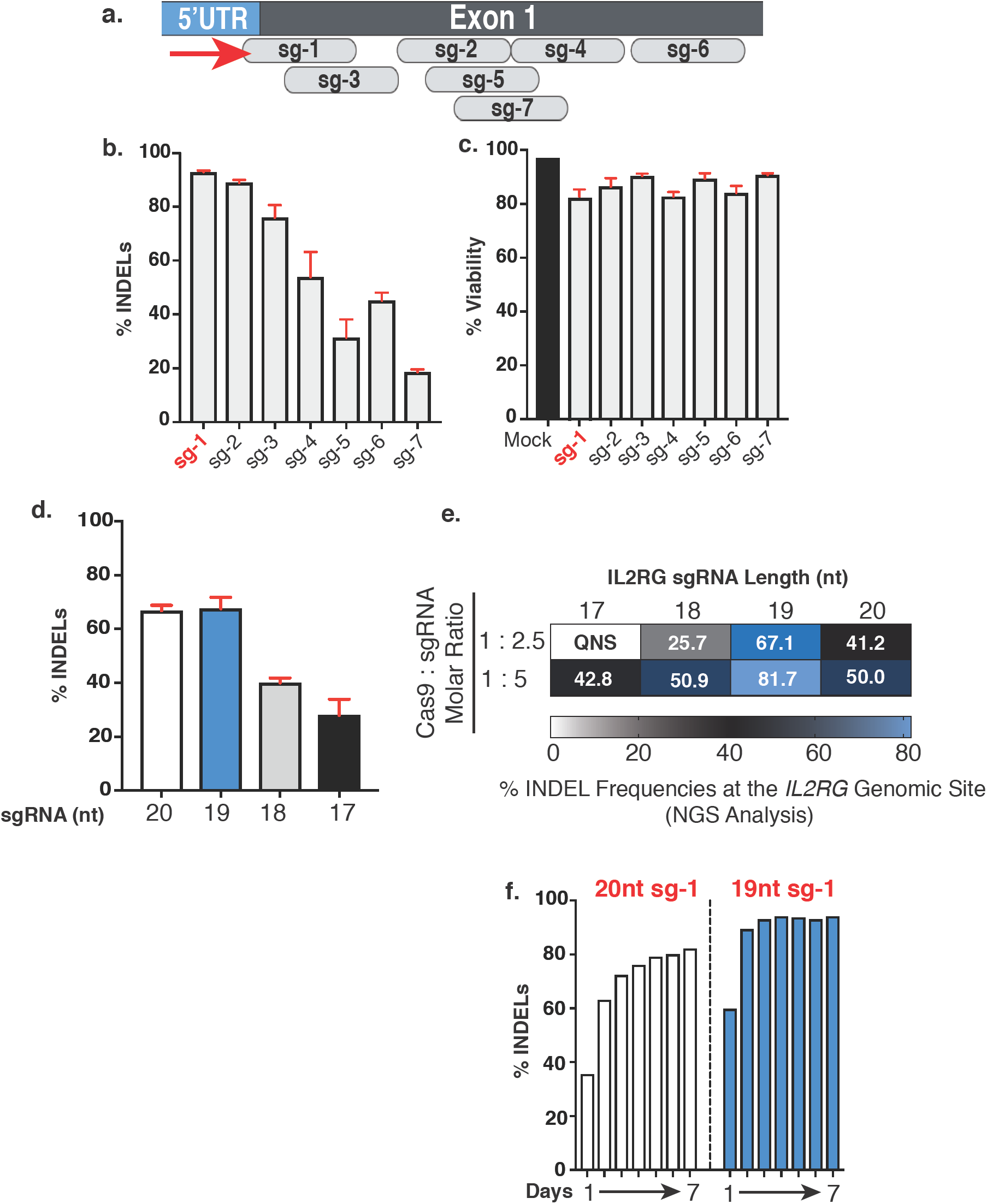
**a.** Schematic of IL2RG sgRNAs for exon 1. **b.** Percent INDELs at day 4 for the seven IL2RG sgRNAs. Percent viability for the sgRNA 1-7 nucleofected as RNP in CB derived CD34^+^ HSPCs. **d.** Comparing percent INDELs of WT (20nt) sg-1 IL2RG sgRNA to truncated versions (19nt, 18nt and 17nt) at 1:25 molar ratio. **e.** Next generation sequencing (NGS) analysis of samples from d. **f.** Time course of percent INDELs generated by WT sg-1 IL2RG sgRNA (white bars) and 19nt truncated IL2Rg sgRNA (blue bars). CB, cord blood.

**Fig. S2.**
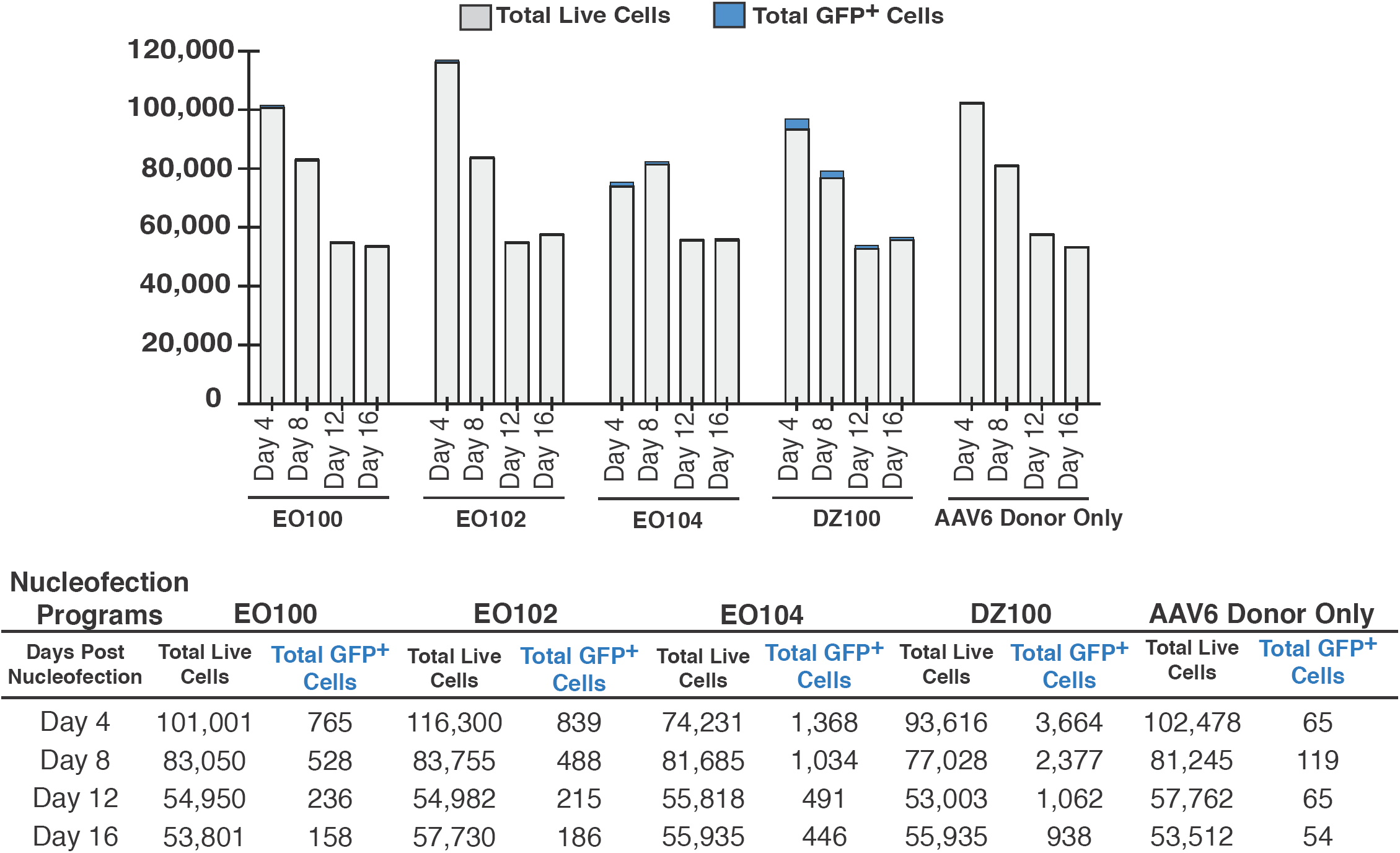
Screening Lonza 4d nucleofection programs for human CD34^+^ HSPCs. 1×10^5^ cord blood derived CD34^+^ HSPCs were nucleofected with RNP at 1: 2.5 molar ratio and transduced with AAV6 donor DNA virus for CCR5 locus at an MOI of 100,000. Total live cells determined by trypan blue staining and % GFP^+^ cells by flow cytometry.

**Fig. S3.**
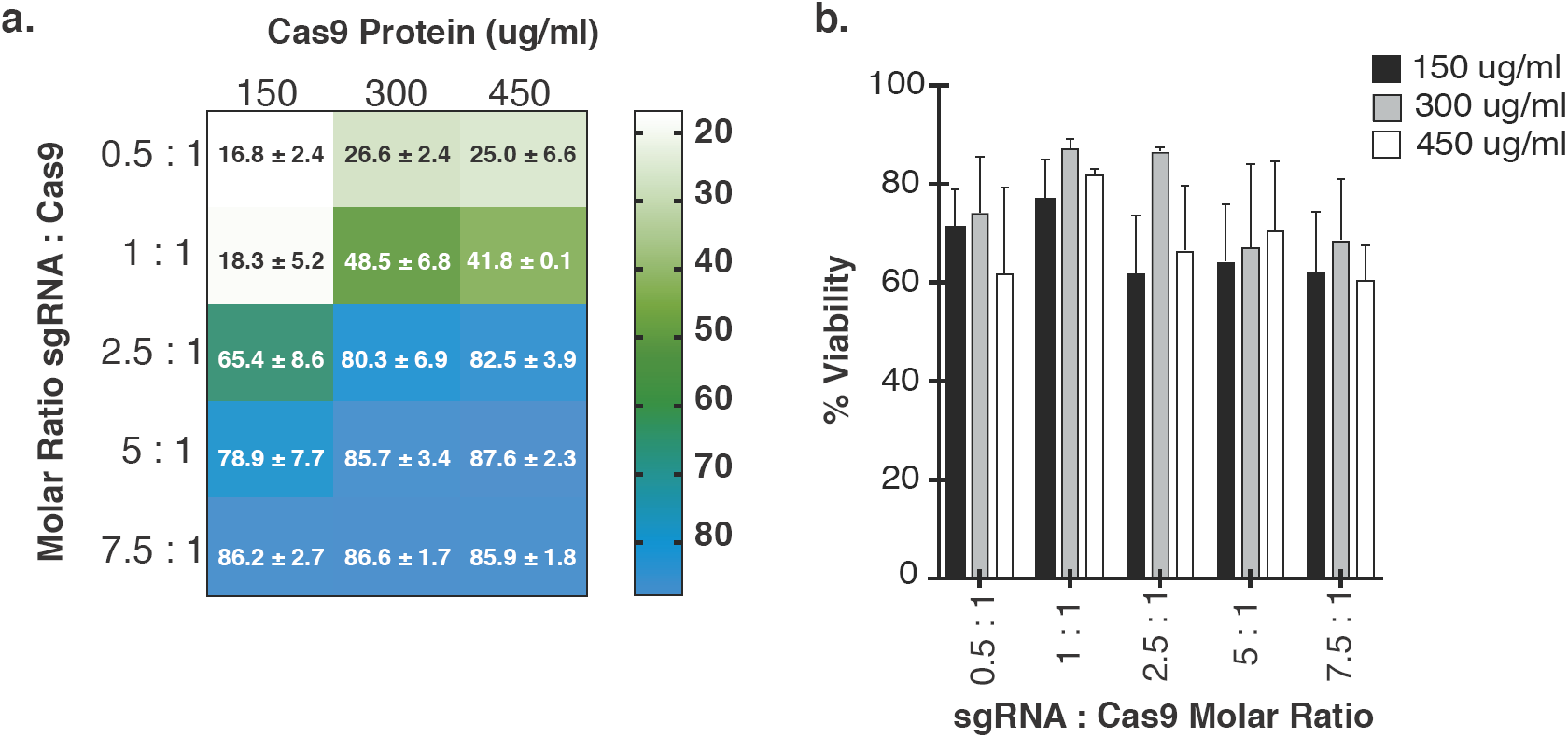
**a.** Molar ratios of IL2RG sgRNA guide and Cas9 protein in mobilized PB CD34^+^ HSPCs. **b.** Heat map TIDE analysis at day 4 post genome editing. RNP complex using *IL2RG* sgRNA sg-1 at various molar ratios. Percent on target INDELs determined from 3 biological replicates (mean ± *s.e.m.*). **b.** Cellular viability at day 4 post nucleofection based on trypan blue stainig. PB, peripheral blood.

**Fig. S4.**
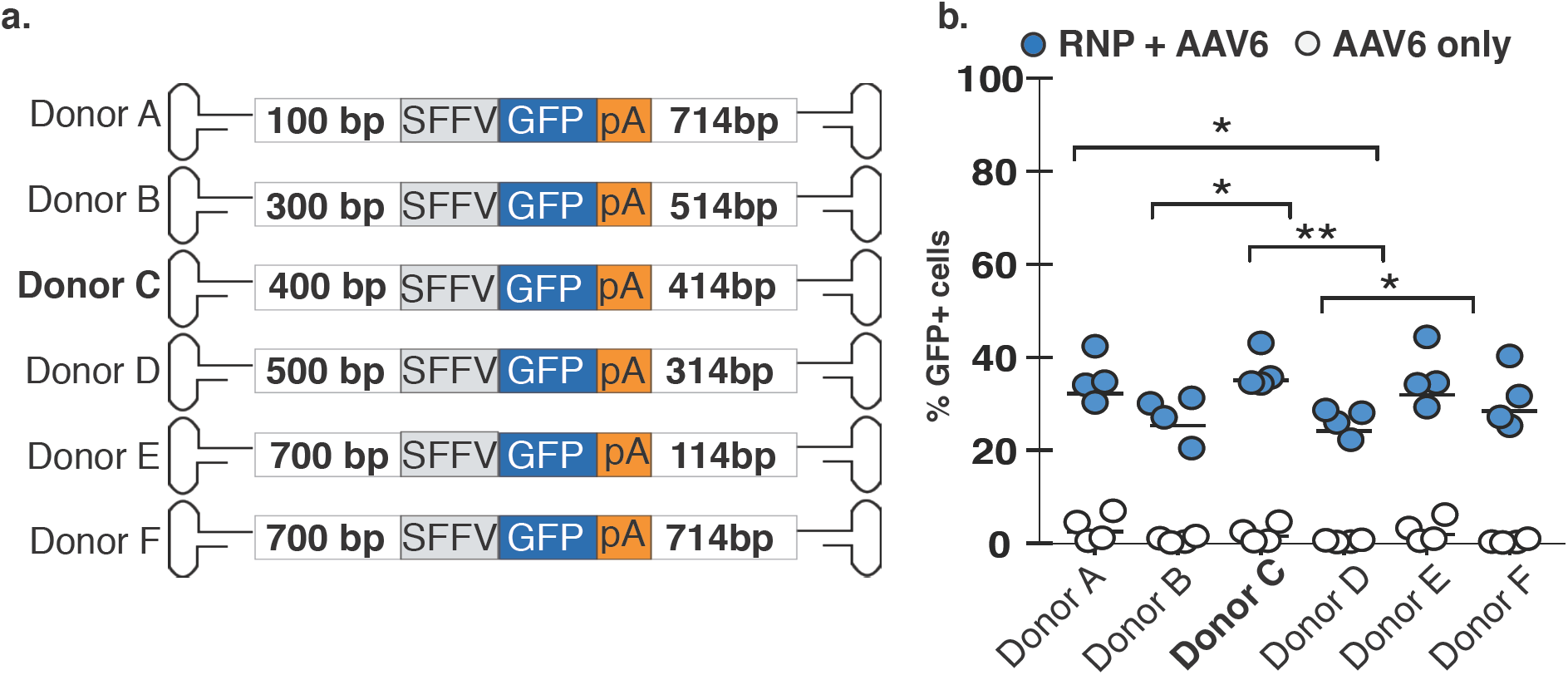
IL2RG specific homology arm lenght screening. **a.** Symmetric and asymmetric arms of homology flanking a SFFV GFP cassette. **b.** Targeting integration frequencies (n= 4 male donors). Donor A vs Donor D *p-value = 0.0204; Donor B vs Donor C *p-value = 0.0226; Donor C vs Donor D **p-value = 0.0055; Donor D vs Donor E *p-value= 0.0361 (unpaired t-test).

**Fig. S5.**
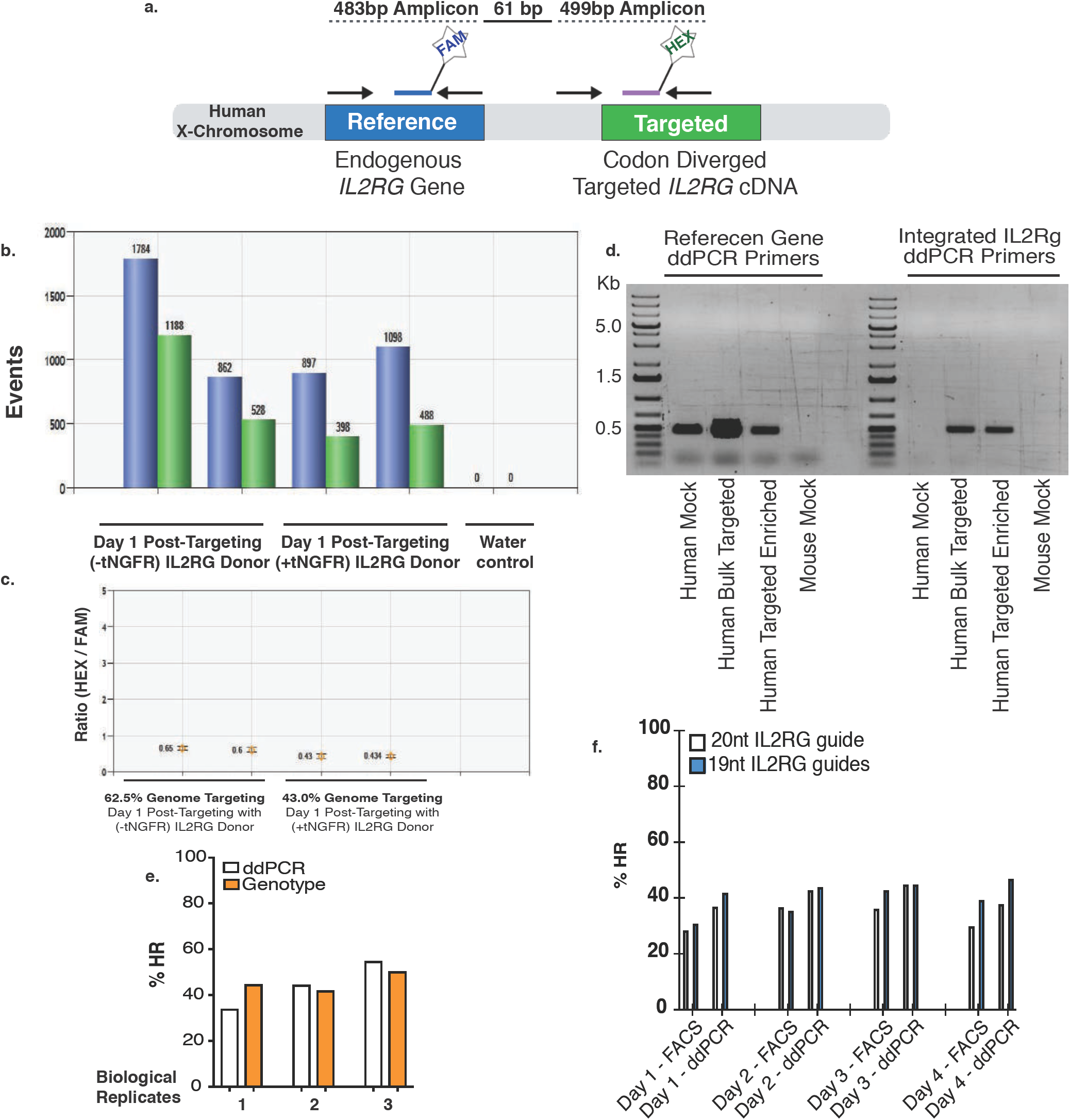
IL2RG specific digital drop PCR (ddPCR) assay. **a.** Schematic representation of the IL2RG specific ddPCR primers/probe design. **b.** Positive droplets generated for the reference (FAM - blue) and integrate (HEX - green) IL2RG PCR amplicons. Genome targeting results using -tNGFR or +tNGFR IL2RG cDNA targeted donor at 24h post rAAV6 transduction. **c.** Ratio of integrated (HEX) over reference (FAM). Male derived genomic DNA contains only one allele of the human X-chromosome allowing for the ratio of the fluorescence signal to be a direct measurement of the levels of genome targeting. **d.** Specificity of the ddPCR primer/probe set. **e.** Comparison of ddPCR analysis of bulk *IL2RG* cDNA targeted male derived CD34^+^ HSPCs and genotype of single cell sorted methylcellulose assay from the bulk population. Shown are three biological replicates. **f.** Comparison of ddPCR and FACS analysis of targeted SFFV-GFP cassette into IL2RG locus of male derived CD34^+^ HSPCs. Time course day 1 through 4 post targeting is shown for full lenght (20nt) and truncated (19nt) IL2RG guide.

**Fig. S6.**
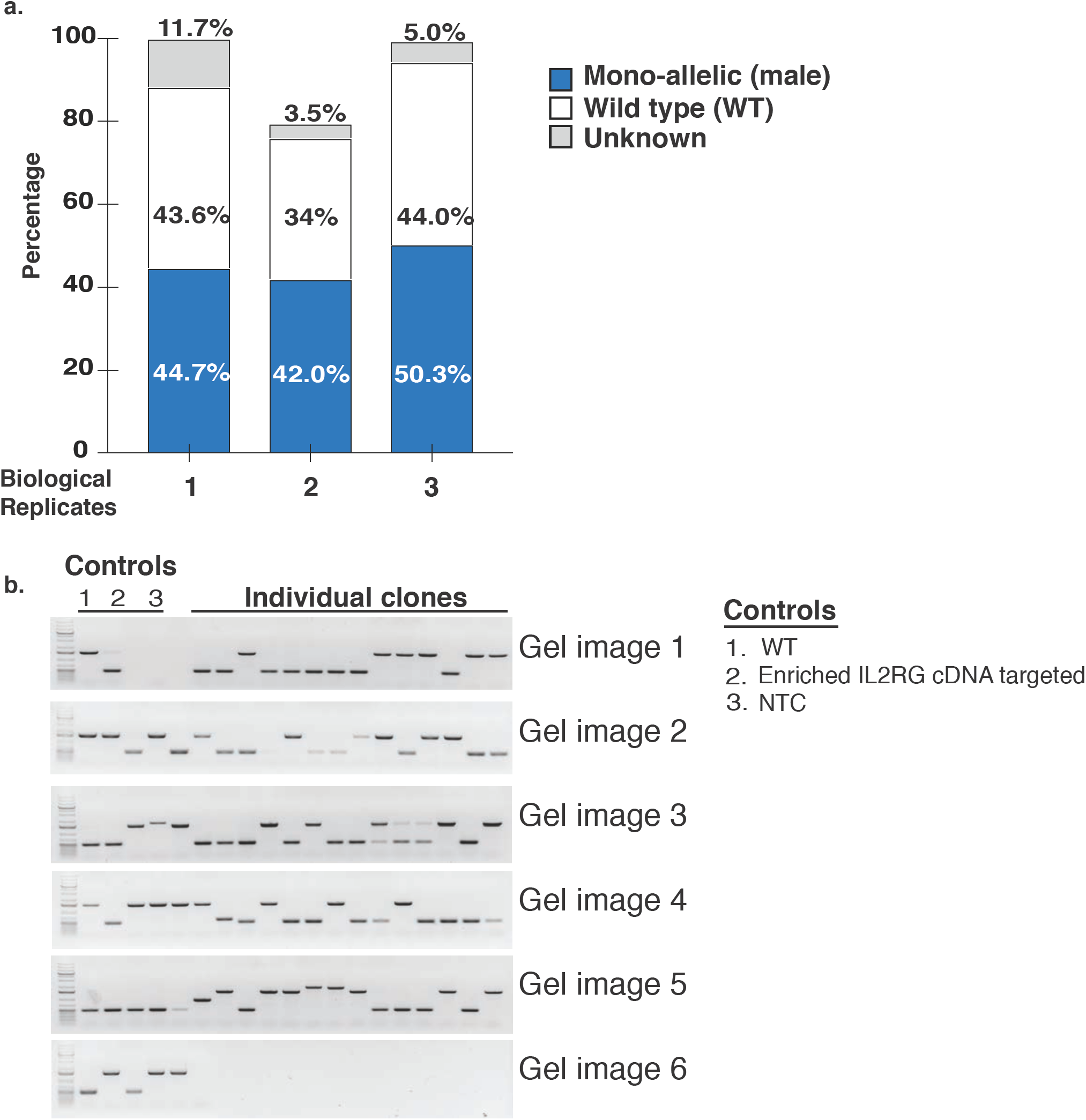
Scoring and genotyping analysis of the IL2RG cDNA targeted, single cell-derived methylcellulose colonies. **a.** Quantification of percent allelic targeting. Genotyping results derived from day 2 post-genome targeting. All three biological replicates were derived from male, frozen mobilized peripheral blood CD34^+^ HSPCs. **b.** Representative genotyping gel images. Colonies obtained at 14 days from individual wells of methylcellulose plates were scored and genotypes using a 3 primer PCR approach. Shown are genotyping results of biological replicate #2. WT - wild type; NTC - no template control.

**Fig. S7.**
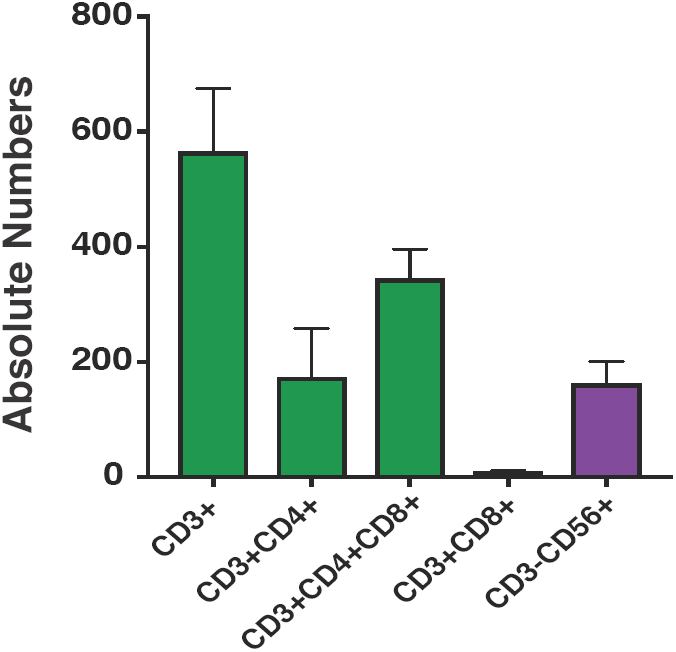
Gene correction of SCID-X1 derived CD34^+^ HSPCs. Shown are absolute numbers of pre-T cells, T-cells and NK cells derived at week one following induction of delta like 1 ligand (dll1) using doxycycline, from Fig. 1e. Bars are average (error bars s.e.m) of 23 wells.

**Fig. S8.**
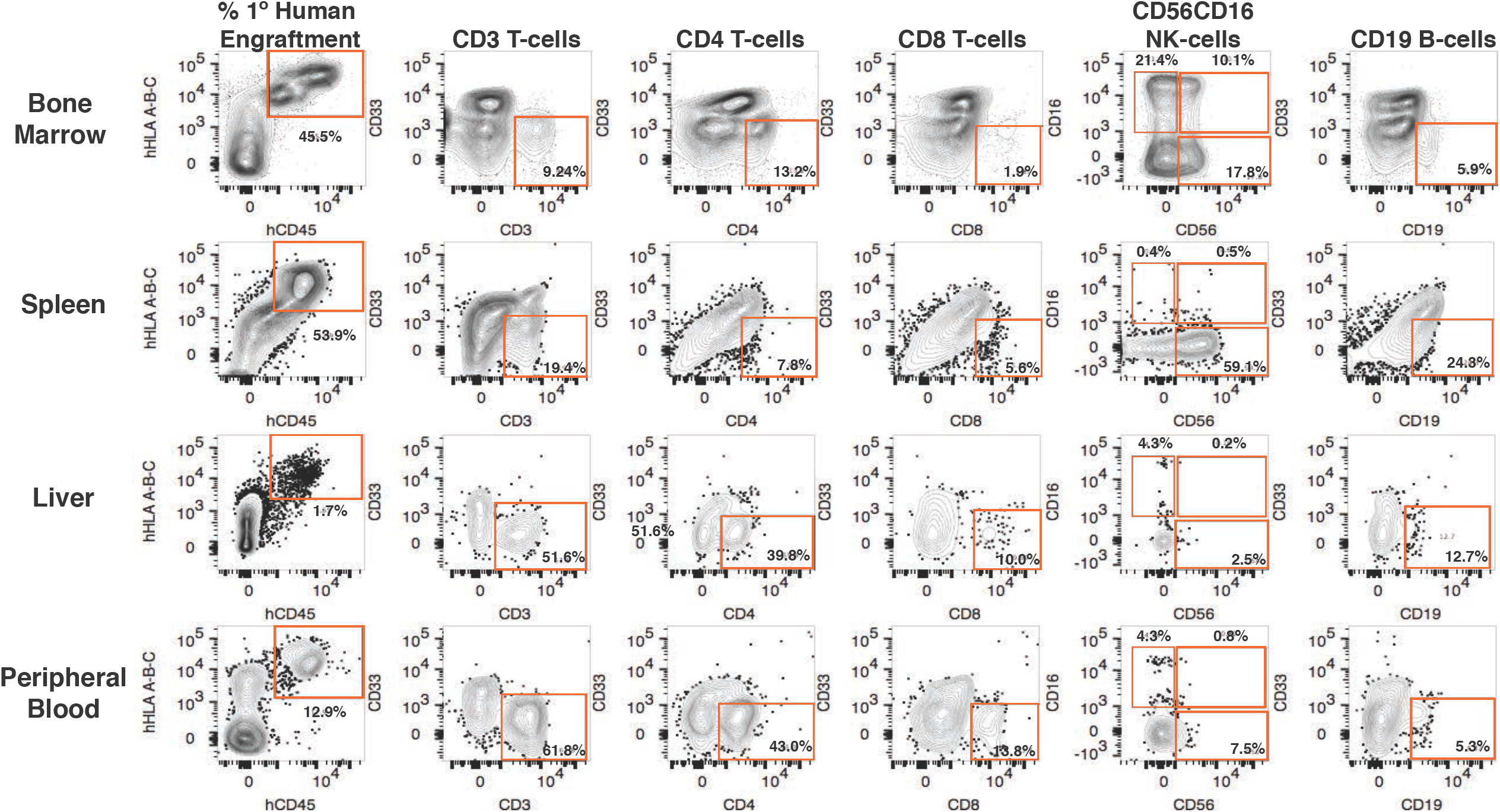
Representative FACS plots for lymphoid lineage analysis at week 16 post intra-hepatic (1°) engraftment of IL2RG targeted CD34^+^ HSPCs into new born NSG mice. Analysis is shown from a mouse with high human engraftment levels.

**Fig. S9.**
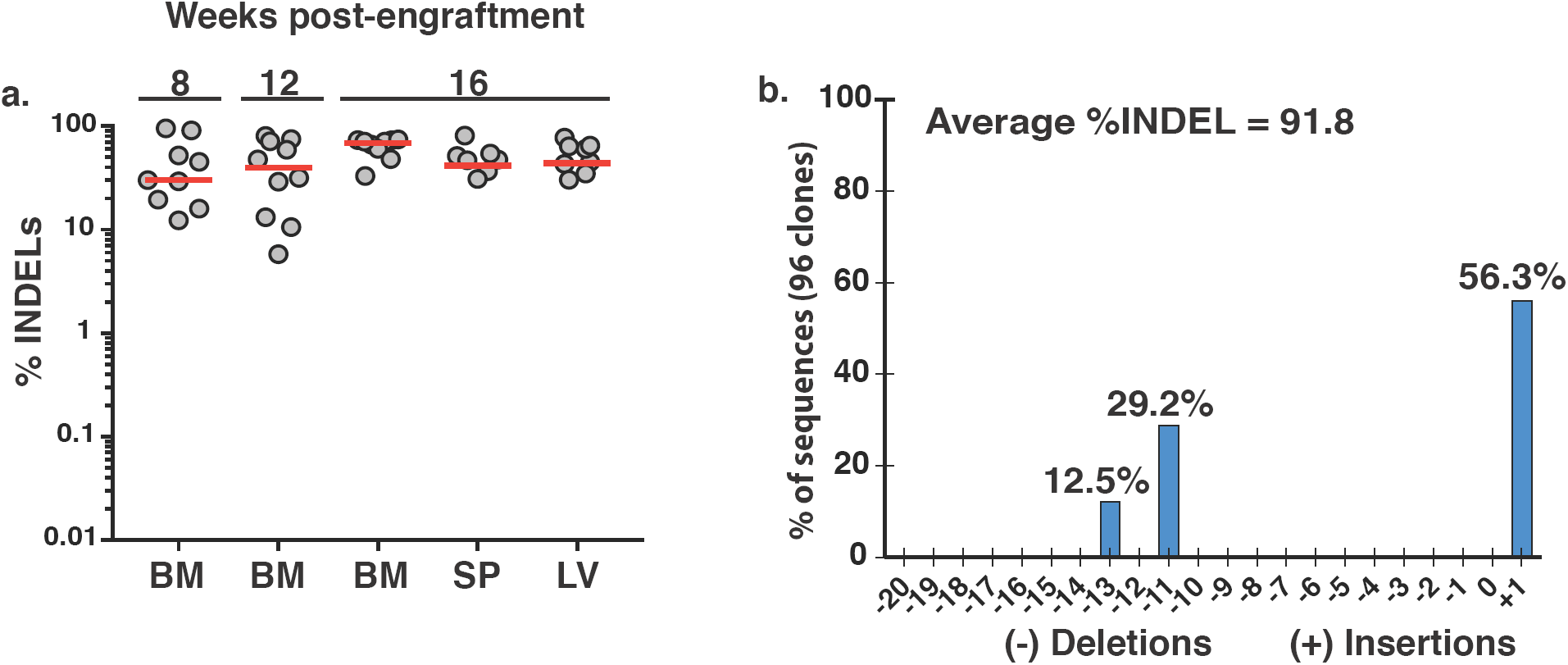
On-target INDEL spectrum analysis of truncated (19nt) IL2RG sgRNA guide#1 in CD34^+^ HSPCs. 1.0×10^5^ CD34^+^ HSPCs derived from male, frozen mobilized peripheral blood (mPB) source, were nucleofected with RNP system at at 5:1 molar ratio. **a.** Percent INDELS determined by TIDE analysis at 8, 12 and 16 weeks post intra-hepatic primary engraftment into NSG pups. **b.** INDEL spectrum characterization generated by truncated 19nt IL2RG guide at day 4 post nucleofection of male derived CD34^+^ HSPCs. Analysis was carried out at clonal level from 96 clones obtained by TOPO clonining bulk RNP sample. IL2RG allele from each 96 clones were sequenced and distribution of INDELs was determined by TIDE analysis.

**Fig. S10.**
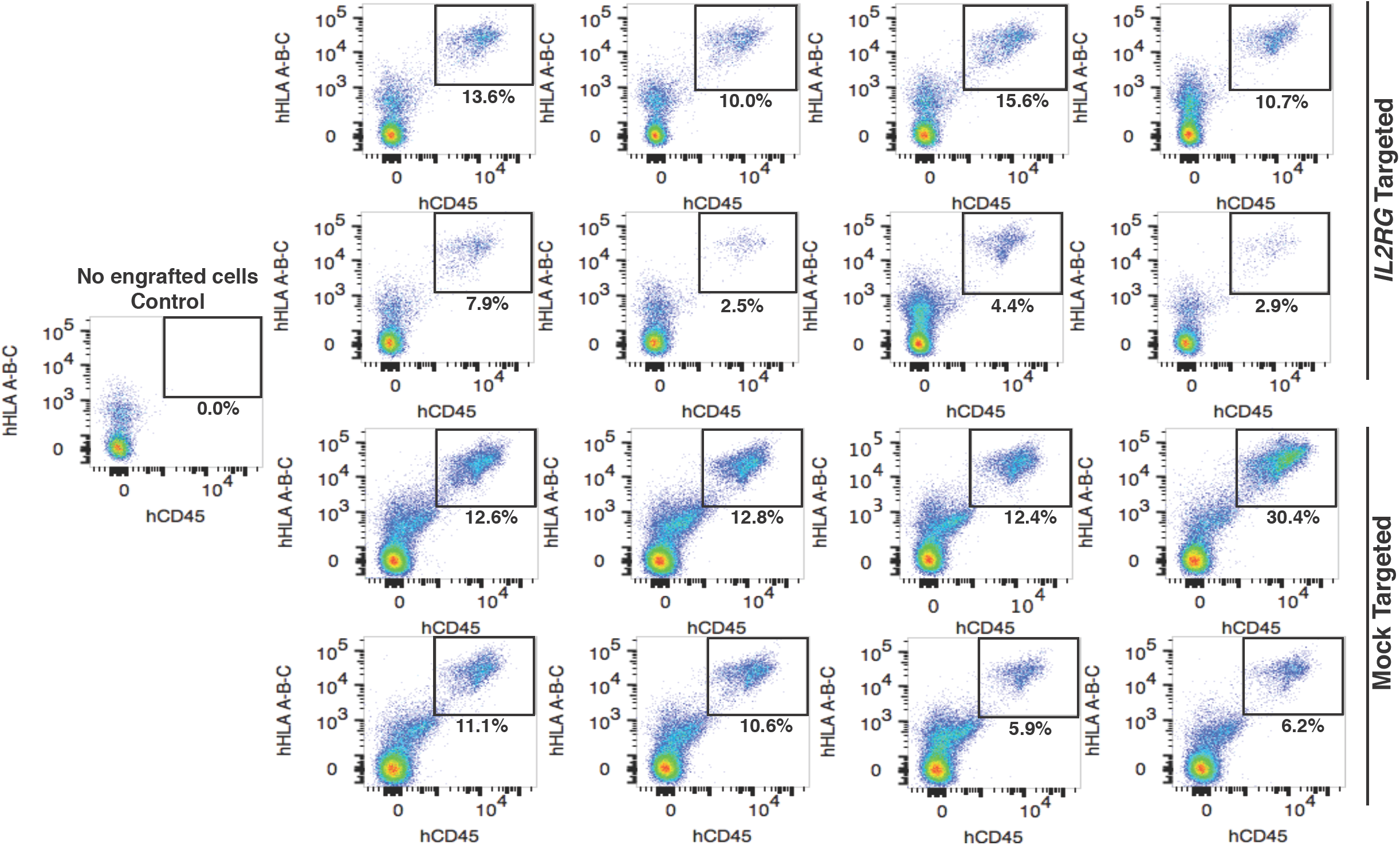
FACS plots showing secondary (2°) human engraftment levels, from total bone marrow, derived from (1°) intra-hepatic IL2RG and mock targeted CD34^+^ HSPCs.

**Fig S11.**
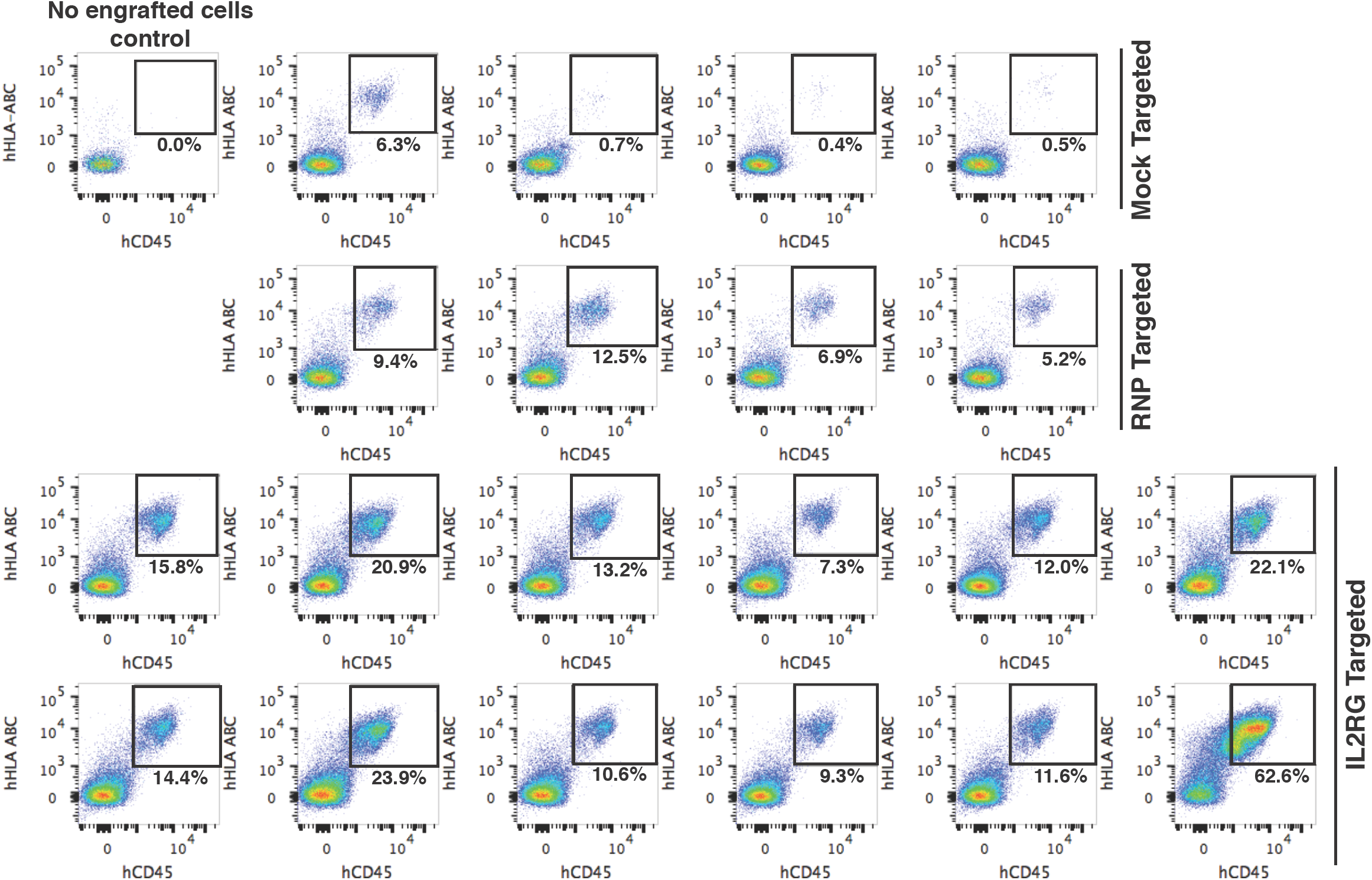
FACS plots showing IF-IF secondary (2°) human engraftment levels in the total bone marrow of mice injected with IL2RG or mock targeted CD34^+^ HSPCs. 5×10^5^ purified CD34^+^HSPCs from total BM of mock or IL2RG targeted engrafted mice were injected IF into sub-lethally irradiated adult NSG mice. The observed low levels of engraftment in 3 out of 4 mice that received mock treated cells were due to fluid backflow during the IF injection procedure. IF: intra-femoral.

**Fig S12.**
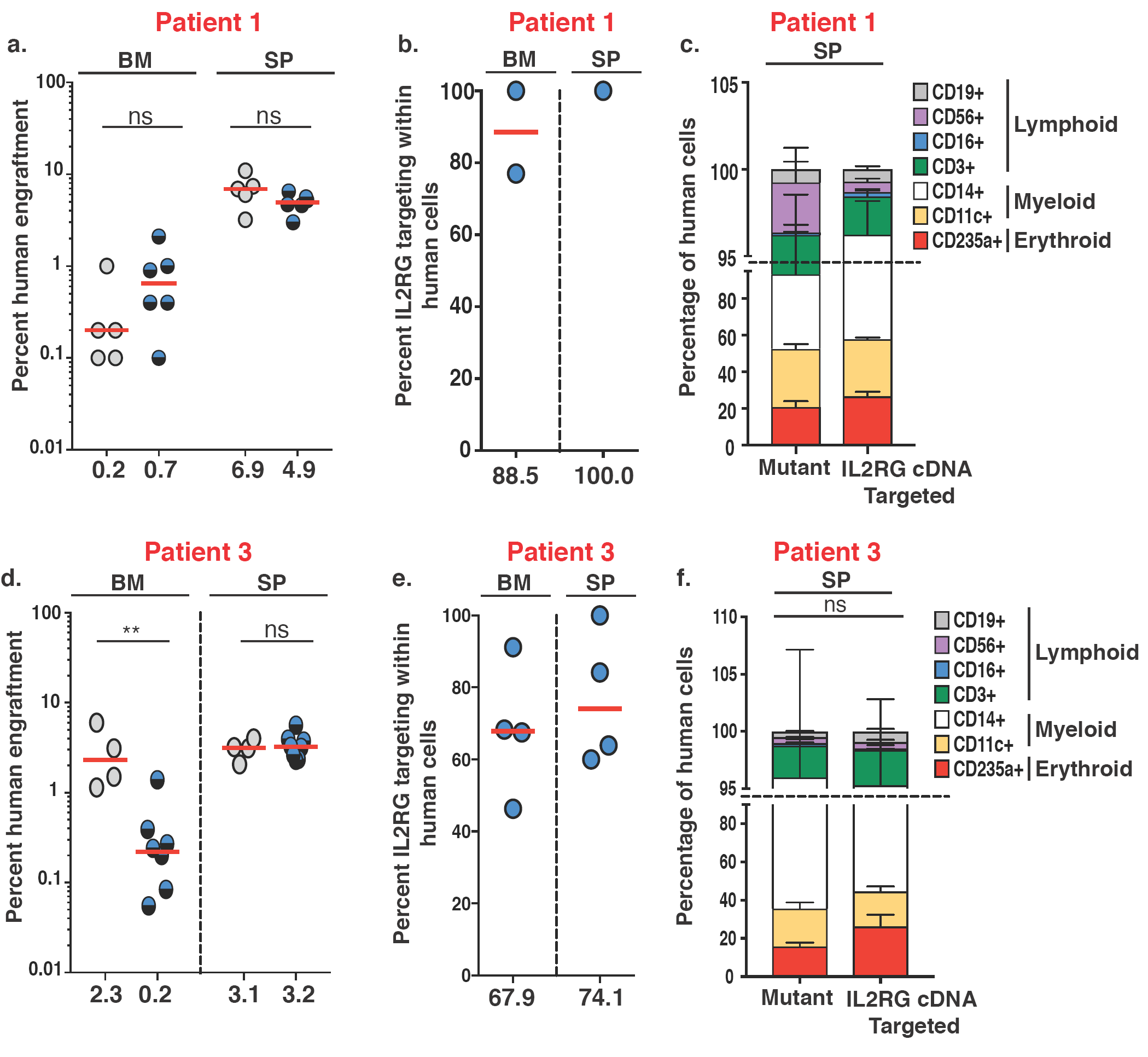
Primary human engraftment of SCID-X1 patient derived HSPCs CD34+. **a.** Human engraftment of mutant or IL2RG targeted HSPCs in bone marrow (BM, n = 5) or spleen (SP, n = 6) 20 weeks after transplant (median plotted). **b.** ddPCR quantification of levels of IL2RG codon-optimized cDNA present in BM (n=2) and SP (n=1) samples. **c.** Percent composition of lymphoid, myeloid and erythroid present in SP 20 weeks post-transplant. **d.** Same as **a**, using SCID-X1 patient 3 derived HSPCs CD34^+^ mutant cells (n = 4) and IL2RG targeted cells (n=7) 17 weeks after transplant (Multiple t-test, Holm-Sidak test, median plotted) **e.** Same as **b,** n = 4**. f.** Same as **c.** \*\**P* = 0.0073, ns, not significant.

**Fig. S13.**
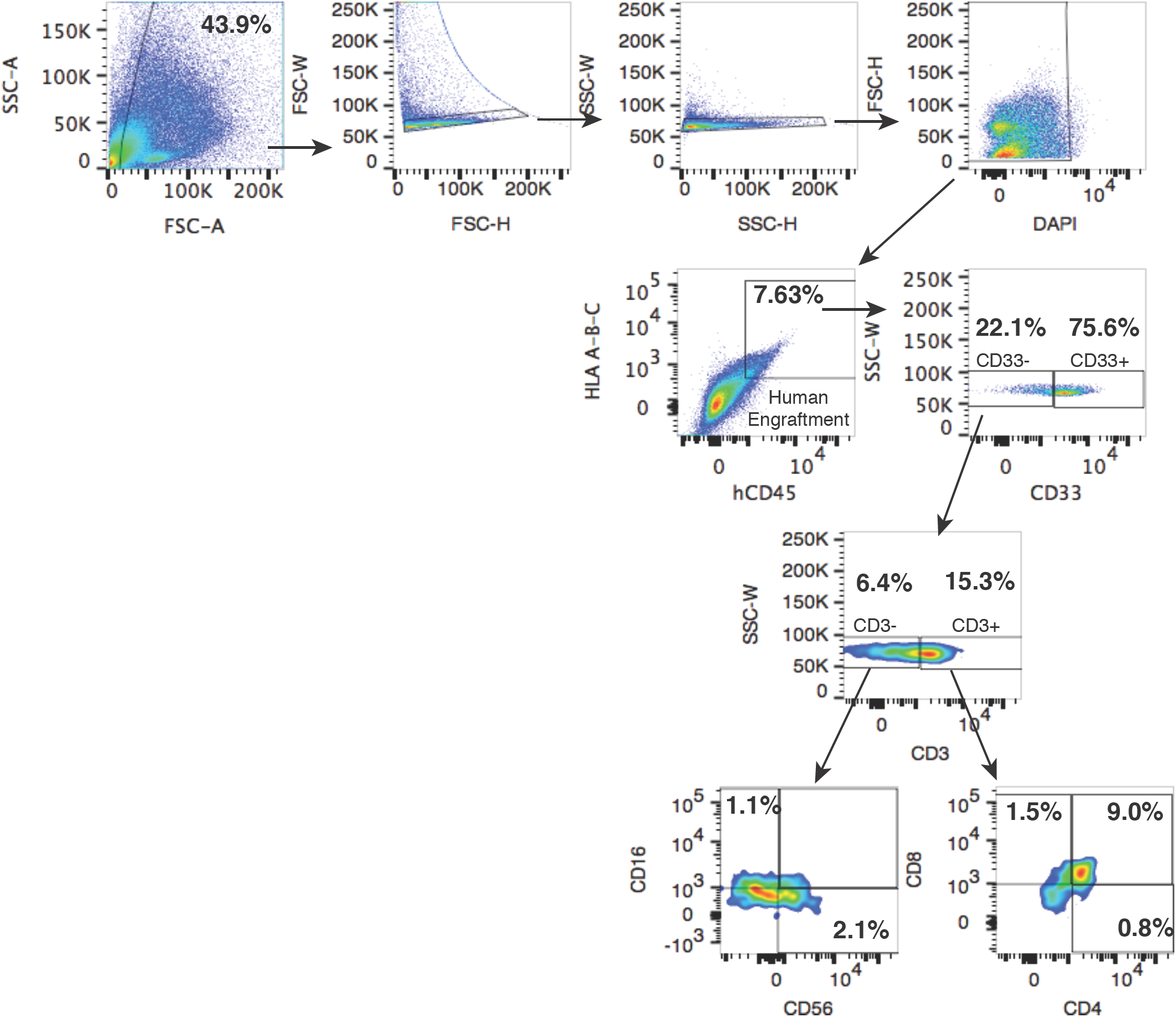
Lymphoid lineage analysis of IL2RG cDNA targeted SCID-X1 patient-derived CD34^+^HSPCs. Representative FACS analysis of spleen sample derived from one NSG mouse at week 16 post-engraftment with IL2RG targeted mobilized PB CD34^+^HSPCs. PB: peripheral blood.

**Fig. S14.**
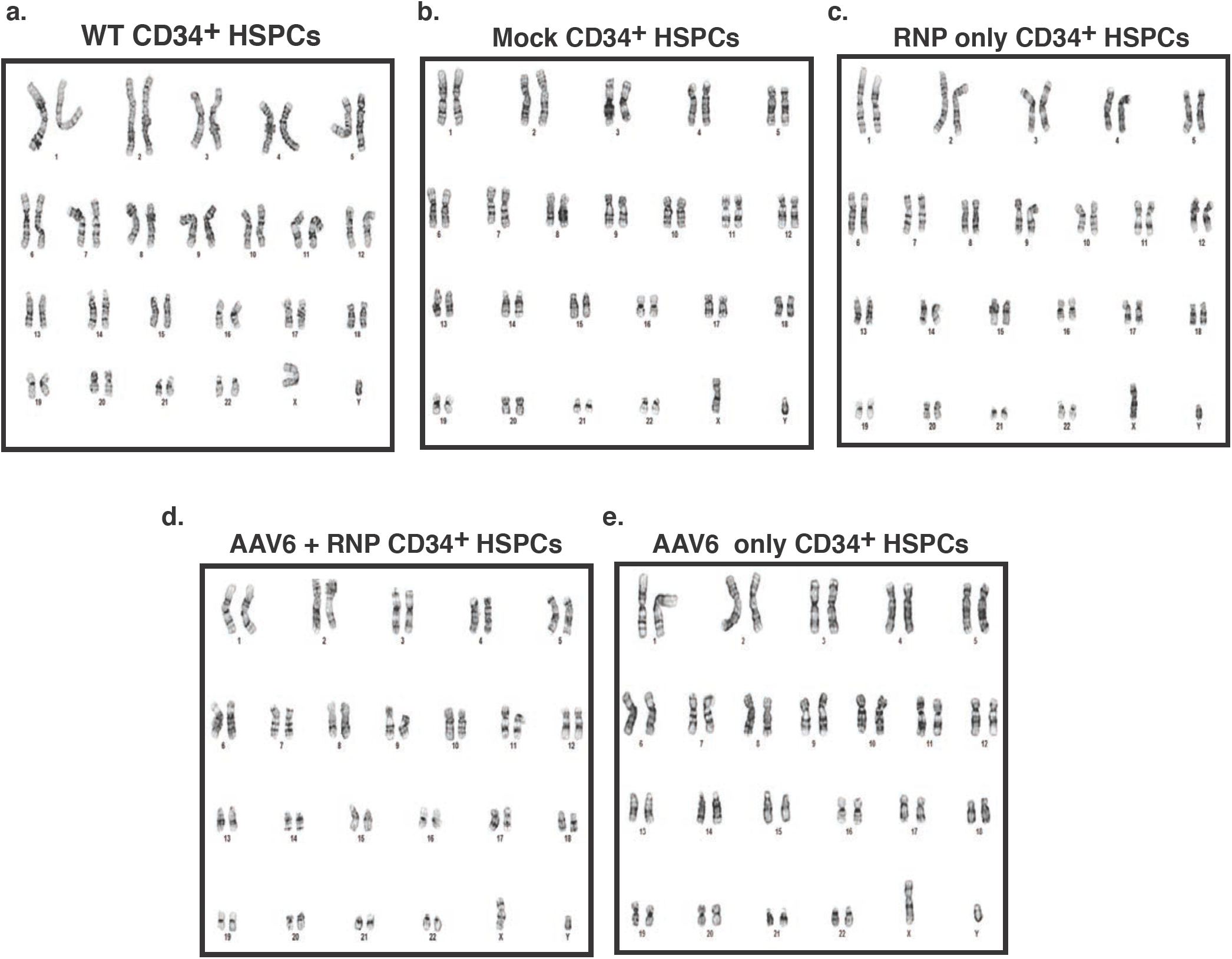
Karyotype analysis of IL2RG genome edited and genome targeted cord blood derived CD34^+^ HSPCs. 5.0×10^5^ cells were nucleofected at day 2 post *ex-vivo* cell culturing with RNP at 5:1 molar ratio. Conditions **(d)** and **(e)** received rAAV6 with -tNGFR IL2RG clinical donor at an MOI of 200,000 vg/ul. Day 2 post transduction, cells were collected and prepared the same day for karyotype analysis. 20 cells were analyzed per condition. Conditions **(c)** and **(d)** and conditions **(d)** and **(e)** show that a combined 40/40 cells treated with RNP or with rAAV6, respectively did not produced cells with chromosomal abnormalities. vg: vector genome.

**Fig. S15.**
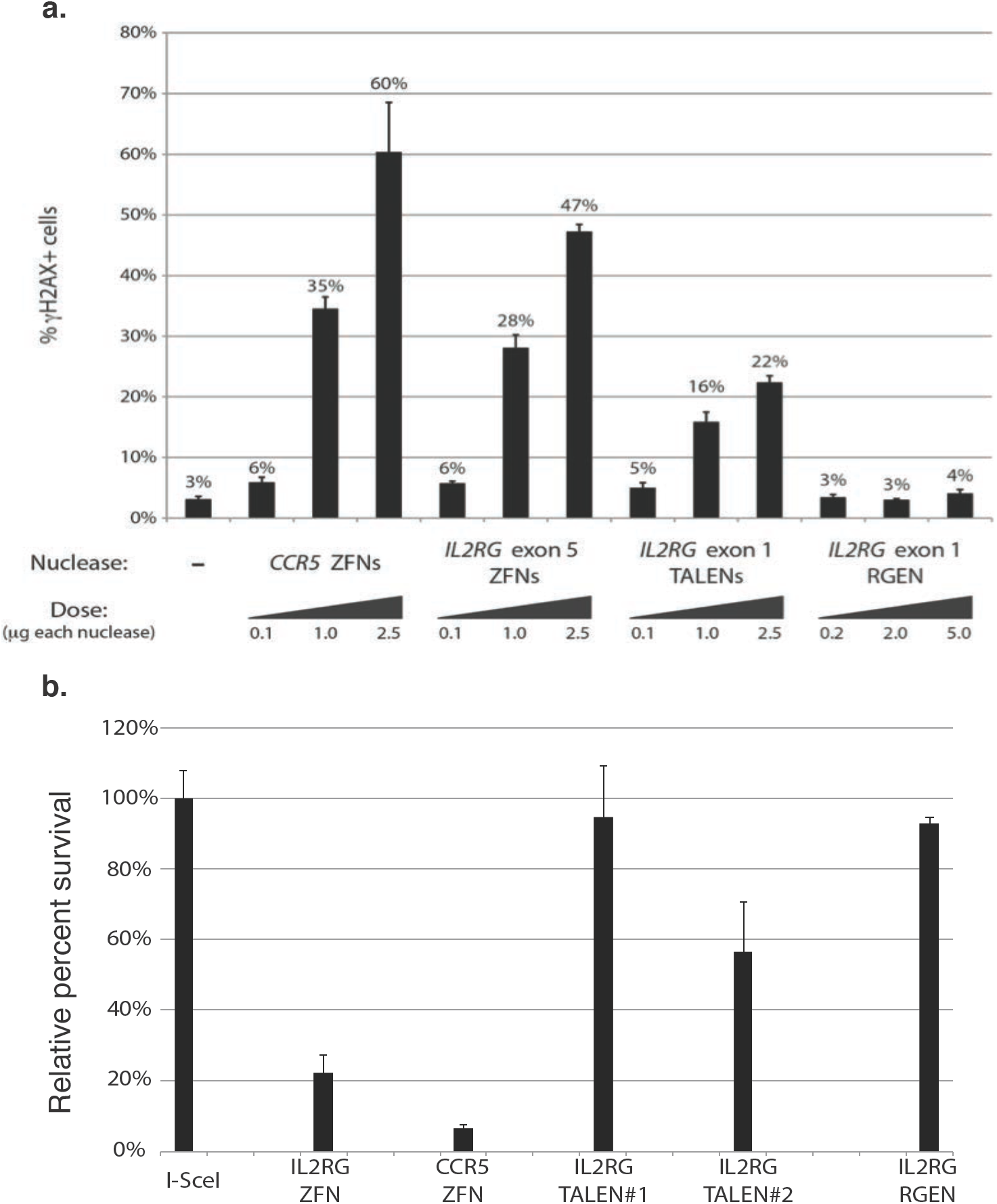
Genotoxicity of IL2RG exon 1 TALENs, IL2RG exon 5 ZFNs, CCR5 ZFNs, and an IL2RG exon 1 RGEN (CRISPR/Cas9). **a.** yH2AX assay. Genotoxicity assay measuring DNA damage induced by different classes of engineered nucleases by assessing the phosphorylation of histone H2AX, a marker of DSB formation, in K562 cells. Percentage yH2AX^+^ cells was measured by flow cytometry 48h post-nucleofection. **b.** Relative cell survival assay. Levels of genotoxicity induced by different classes of engineered nucleases in 293T cell line. Cells were nucleofected with GFP plasmid DNA and genome wide off-target activity of each nuclease was determined by FACS analysis as percent GFP^+^ cells relative to I-SceI control. Bars (n = 3, mean ± s.d)

